# Network instability dynamics drive a transient bursting period in the developing hippocampus *in vivo*

**DOI:** 10.1101/2021.05.28.446133

**Authors:** Jürgen Graf, Vahid Rahmati, Myrtill Majoros, Otto W. Witte, Christian Geis, Stefan J. Kiebel, Knut Holthoff, Knut Kirmse

## Abstract

Spontaneous correlated activity is a universal hallmark of immature neural circuits. However, the cellular dynamics and intrinsic mechanisms underlying network burstiness in the intact developing brain are largely unknown. Here, we use two-photon Ca^2+^ imaging to comprehensively map the developmental trajectories of spontaneous network activity in hippocampal area CA1 *in vivo*. We unexpectedly find that network burstiness peaks after the developmental emergence of effective synaptic inhibition in the second postnatal week. We demonstrate that the enhanced network burstiness reflects an increased functional coupling of individual neurons to local population activity. However, pairwise neuronal correlations are low, and network bursts (NBs) recruit CA1 pyramidal cells in a virtually random manner. Using a dynamic systems modeling approach, we reconcile these experimental findings and identify network bi-stability as a potential regime underlying network burstiness at this age. Our analyses reveal an important role of synaptic input characteristics and network instability dynamics for NB generation. Collectively, our data suggest a mechanism, whereby developing CA1 performs extensive input-discrimination learning prior to the onset of environmental exploration.

## Introduction

Developing neural circuits generate correlated spontaneous activity in which co-activations of large groups of neurons are interspersed by relatively long periods of quiescence (Kirmse et al., 2022; Molnar et al., 2020). In rodents, network activity commences long before the onset of hearing, vision and active environmental exploration and makes important contributions to the proper assembly of brain circuits (Kirkby et al., 2013). Activity-dependent refinements operate at multiple steps of maturation, including the control of neural progenitor progression (Vitali et al., 2018), apoptotic cell death (Blanquie et al., 2017; Wong et al., 2018), neuronal cell-type specification (Sun et al., 2018), migration (Maset et al., 2021) as well as synapse formation and plasticity (Oh et al., 2016; Sando et al., 2017; Winnubst et al., 2015). Experimental and theoretical evidence suggests that, in addition to the overall level of activity, specific spatiotemporal firing patterns are critical for activity-dependent refinements to occur (Albert et al., 2008; Zhang et al., 2011).

A representative example of correlated spontaneous network activity is found in the neonatal hippocampus *in vivo*. During the first postnatal week, the main electrophysiological signature are bursts of multi-unit activity (MUA) (Leinekugel et al., 2002), which bilaterally synchronize large parts of the dorsal CA1 and are often accompanied by sharp waves (SPWs) in the local field potential (Valeeva et al., 2019; Valeeva et al., 2020). SPWs frequently follow myoclonic limb or whisker twitches (Dard et al., 2022; Del Rio-Bermudez et al., 2020; Karlsson et al., 2006; Valeeva et al., 2019), suggesting that SPWs convey feedback information from the somatosensory periphery. By the second postnatal week, discontinuous activity in the olfactory bulb drives network oscillations in the entorhinal cortex (Gretenkord et al., 2019), further pointing to a role of multi-sensory integration in limbic ontogenesis. In the neonatal CA1, recent *in vivo* investigations revealed that GABAergic interneurons (INs) could promote a second class of SPW-independent network events through NKCC1-dependent chloride uptake in pyramidal cells (PCs), although inhibitory effects of GABAergic signaling coexist (Graf et al., 2021; Murata et al., 2020; but see Valeeva et al., 2016). A qualitatively similar situation applies to the immature hippocampus *in vitro* (Ben-Ari et al., 1989; Flossmann et al., 2019), in which correlated spiking of PCs is facilitated by increasing the intracellular chloride concentration (Spoljaric et al., 2019; Zhang et al., 2019), whereas inhibition of chloride uptake has the opposite effect (Dzhala et al., 2005). In this line, *in vitro* studies suggest that correlated spontaneous activity largely disappears by the beginning of the second postnatal week, when the reversal potential of GABA_A_ receptor-mediated currents shifts into the hyperpolarizing direction (Spoljaric et al., 2017; Tyzio et al., 2008). However, the developmental trajectories of cellular network firing dynamics in the hippocampus *in vivo* remain largely unknown.

Using two-photon Ca^2+^ imaging, we here provide a detailed analysis of the spatiotemporal dynamics of network activity in the developing CA1 region at single-cellular resolution *in vivo*. We reveal that CA1 PCs undergo a transient period of enhanced burst-like network activity during the second postnatal week, when GABA already acts as an inhibitory transmitter. Our results show that, at this time, network bursts (NBs) recruit CA1 PCs in an almost random manner, and recurring cellular activation patterns become more stable only after eye opening. Using computational network modeling, we identify bi-stability as a dynamical regime underpinning the enhanced bursting activity of CA1 PCs. We show that NBs mainly reflect the network’s intrinsic instability dynamics, which exquisitely depend on proper input timing and strength. In addition, inhibitory GABAergic signaling effectively promotes state transitions underlying NB generation. Our data suggest a mechanism, whereby CA1 undergoes extensive input-discrimination learning before the onset of environmental exploration.

## Results

### Reliable detection of somatic Ca^2+^ transients in densely labeled tissue

We used *in vivo* two-photon laser-scanning microscopy (2PLSM) in spontaneously breathing, head-fixed *Emx1^IREScre^:GCaMP6s^LSL^* mice to record somatic Ca^2+^ transients (CaTs) from CA1 PCs as a proxy of their firing activities. In this strain, Cre is expressed in virtually all CA1 PCs (Gorski et al., 2002; Kummer et al., 2012). Due to the finite point-spread function inherent to 2PLSM, dense cell labeling resulted in a non-negligible overlap of signals originating from neighboring somata and/or neurites (Chen et al., 2020; Denis et al., 2020). Our preliminary analysis revealed that, under such conditions, standard CaT detection methods based on analyzing mean fluorescence intensities from regions of interests (ROIs) can lead to substantial false positive rates (Fig. 1). We therefore devised a novel cell-specific spatial template-matching approach for the reliable detection of CaTs in densely labeled tissue, which we refer to as CATHARSiS (<u>Ca</u>lcium transient detection <u>har</u>nessing <u>s</u>patial <u>s</u>imilarity). CATHARSiS makes use of the fact that, for each cell, the spike-induced changes in GCaMP fluorescence intensity (ΔF) have a spatially inhomogeneous (ring-like) configuration (see Methods for details). In brief, a cell-specific spatial ΔF template representing the active cell is computed (Figs. 1A and 1B) and optimally scaled to fit its ΔF in each recorded frame. Based on the optimum scaling factor and the quality of the fit, a detection criterion *D(t)* is computed for each time point (Clements et al., 1997). *D(t)* is then subjected to a general-purpose event detection routine for the extraction of CaT onsets (Rahmati et al., 2018). We first illustrate CATHARSiS by analyzing simulated spike-induced CaTs in ring-shaped cells (Figs. 1A and 1B). Here, fluorescence signals of the cell of interest were contaminated by (I) signals originating from a partially overlapping second cell, (II) spatially homogenous fluorescence changes mimicking axon-based neuropil activity (Kerr et al., 2005) and (III) a low level of Poissonian noise (Fig. 1C). Figures 1C and 1D demonstrate that *D(t)* will increase only if ΔF has a spatial configuration similar to that of the template, i.e. if the simulated cell is active. Of note, *D(t)* is insensitive to a spatially uniform offset of ΔF and can decrease for mean ΔF increases having a dissimilar spatial configuration (#2 and #3 in Figs. 1C and 1D). We applied CATHARSiS to two simulated sample cells of identical shape and varied their spatial overlap from 0 to 75% of the cell area, in accordance to the observed overlap in our empirical data. CATHARSiS correctly retrieved all ground-truth CaTs without false positive events (n = 665 CaTs in total). We also found that the delay of detected CaT onsets *vs.* simulated spikes was low (−0.3 ± 0.0 frames), pointing to a high temporal accuracy of spike reconstruction, which is a prerequisite of a precise analysis of spatiotemporal activity patterns. We next evaluated CATHARSiS on data recorded from developing CA1 PCs in *Emx1^IREScre^:GCaMP6s^LSL^* mice *in vivo* (Figs. 1E–I). For comparison, a consensus visual annotation by human experts was used (Fig. 1G, top), as simultaneous electrophysiological data were not available (see Methods). We compared CATHARSiS to an event detection routine based on analyzing mean ΔF(t) and found that recall was ∼95% for both approaches (Fig. 1I; see Table S1). However, CATHARSiS yielded considerably fewer false positive events, thus resulting in a significantly higher precision and F1 score (Fig. 1I). Importantly, the delay of detected CaT onsets relative to the consensus annotation was consistently low (0.9 ± 0.1 frames at a frame rate of 11.6 Hz, n = 20 cells), confirming that CATHARSiS achieved a high temporal accuracy.

**Figure 1.**
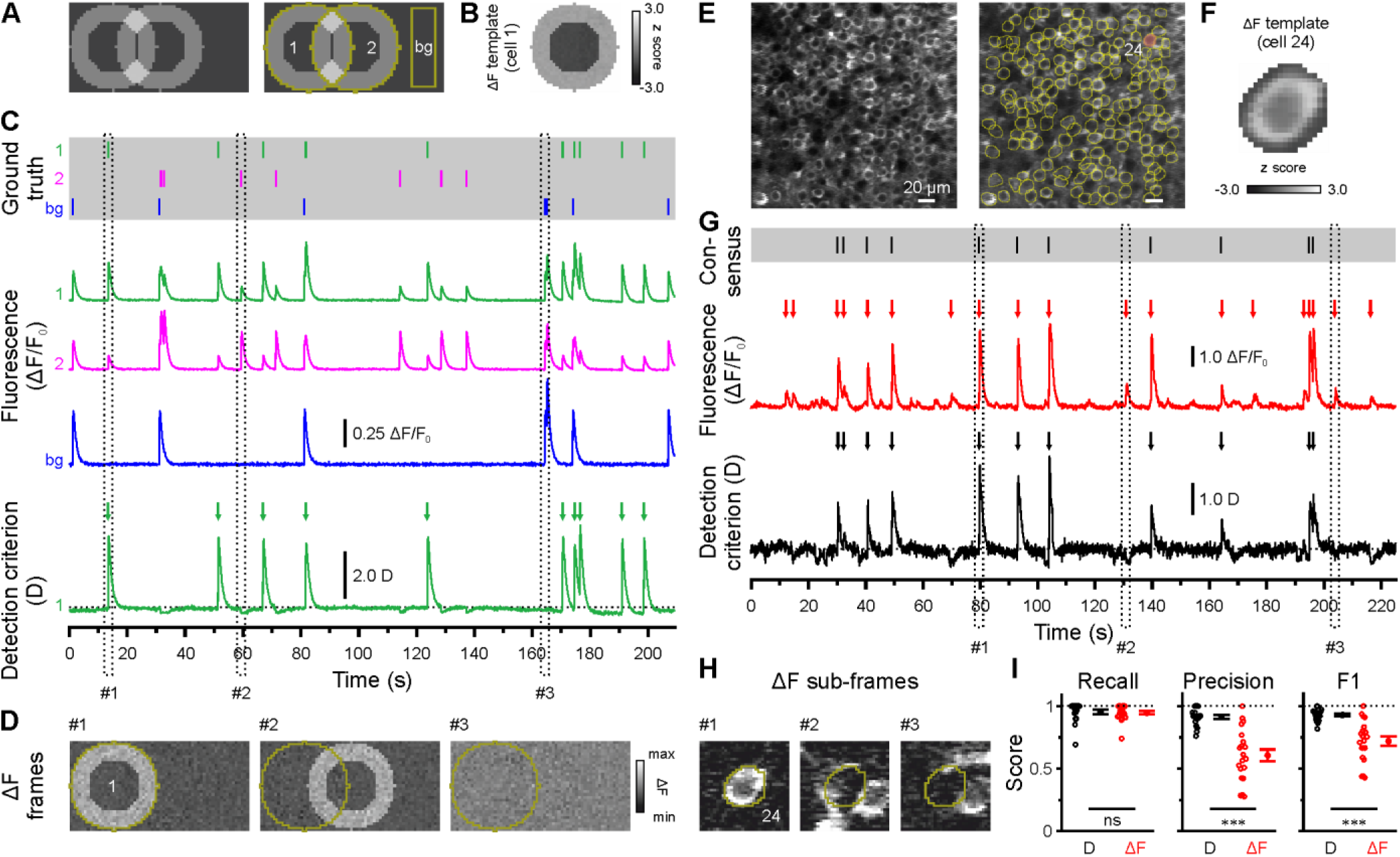
CATHARSiS enables reliable CaT detection in densely labeled tissue. (A) Resting image of two partially overlapping simulated cells (*left*) and regions of interest (ROIs) used for analysis (*right*). bg - background. (B) ΔF template of cell 1. (C) *Top*, simulated trains of action potentials. *Middle*, relative changes from baseline fluorescence (ΔF/F_0_) of ROIs shown in A. *Bottom*, detection criterion (D) for cell 1 and corresponding CaT onsets retrieved by CATHARSiS (arrows). (D) Sample ΔF images of three individual frames at time points indicated in C. Spikes in cell 1 or 2 translated into ring-shaped increases in ΔF, whereas those induced by bg spikes were applied to the entire field of view. (E) Resting GCaMP6s fluorescence image (*left*) and ROIs used for analysis (*right*). (F) ΔF template of the cell indicated in E. (G) *Top*, consensus visual annotation by two human experts for the same cell. *Middle*, ΔF/F_0_ and detected event onsets (red arrows). *Bottom*, detection criterion (D) and corresponding CaT onsets retrieved by CATHARSiS (black arrows). (H) Sample ΔF images of three individual frames at time points indicated in G. Note that frames #2 and #3 led to false positive results if event detection was performed on mean ΔF, but not if performed on D. (I) Quantification of recall, precision and F1 score for event detection based on D (i.e. CATHARSiS) and mean ΔF, respectively. Each open circle represents a single cell. Data are presented as mean ± SEM. ns – not significant. ^***^ P < 0.001. See also Table S1 and Source Data File.

We conclude that CATHARSiS is suited for the reliable reconstruction of somatic CaTs in densely packed neuronal tissue with both high detection and temporal accuracies.

### A transient period of firing equalization during CA1 development in vivo

In the adult CA1, firing rate distributions are approximately log-normal, implying that a minority of neurons accounts for the majority of spikes. In addition, firing rates of individual cells are relatively stable across brain states and tasks, suggesting that skewed firing rate distributions reflect an inherent characteristic of mature hippocampal computations (Mizuseki et al., 2013). To reveal developmental trajectories of single-cell activity, we applied CATHARSiS to extract spontaneous CaTs from *Emx1*+ PCs at P3–4 (n = 19 fields of view [FOVs] from six mice), P10– 12 (n = 11 FOVs from six mice) and P17–19 (n = 12 FOVs from six mice) (Fig. 2A). For the sake of brevity, these age groups are hereafter referred to as P4, P11 and P18, respectively. We found that mean CaT frequencies significantly increased ∼2.5-fold from 1.5 ± 0.2 min^-1^ at P4 to 3.9 ± 0.4 min^-1^ at P11 and remained relatively stable afterwards (P18: 4.8 ± 0.3 min^-1^, Figs. 2B and 2C; see Table S2). Additionally, we observed a striking change in the shape of CaT frequency distributions, which were broad and strongly right-tailed at P4 and P18, but much less so in the second postnatal week (Fig. 2B). To quantify the dispersion of CaT frequencies among individual cells, we plotted the corresponding Lorenz curves (Fig. 2D), in which the cumulative proportion of CaT frequencies is plotted against the cumulative proportion of cells rank-ordered by frequency (Mizuseki et al., 2013). Here, the line of equality represents the case where all neurons have equal CaT frequencies. We computed the Gini coefficient as a measure of deviation from equality (for a graphical representation, see inset in Fig. 2D). Gini coefficients underwent a transient minimum at P11, indicating that CaT frequencies among individual neurons were considerably more similar to each other as compared to P4 and P18 (Fig. 2E). We next addressed whether developmental alterations in average CaT frequencies were accompanied by changes in their irregularity of occurrence. The local coefficient of variation (CV2), a robust measure of local spiking irregularity (Holt et al., 1996; Ponce-Alvarez et al., 2010), gradually declined from P4 to P18 (Fig. 2G). At P11, CV2 was close to one, indicating that the irregularity of CaT occurrence is similar to that of a Poissonian point process, in which successive events occur independently of one another. As previously observed for CaT frequencies, CV2 distributions were also relatively broad at P4 and P18, but narrow at P11 (Fig. 2F). Consistently, Gini coefficients of CV2 showed a distinct minimum at P11 (see #4 in Table S2).

**Figure 2.**
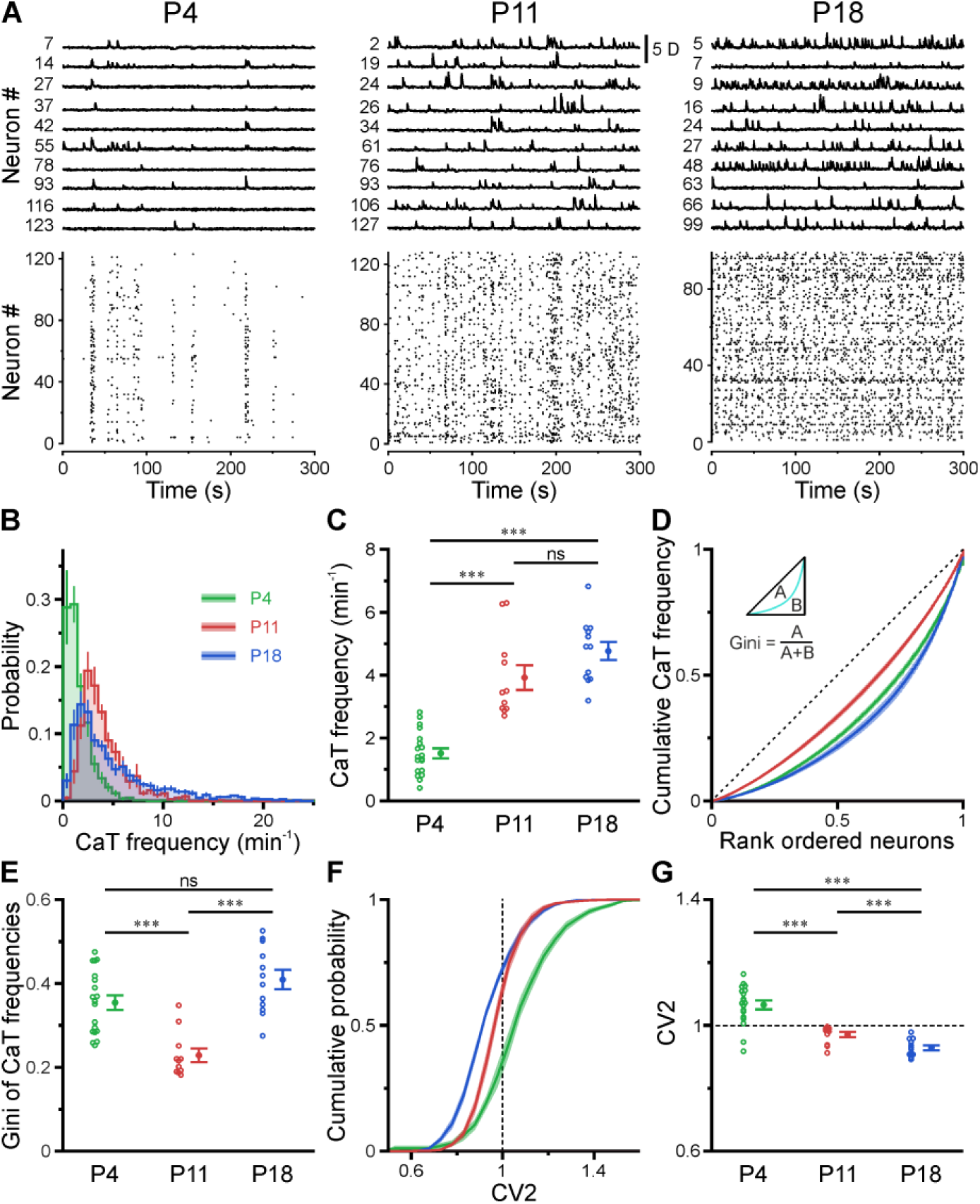
A transient period of firing equalization during CA1 development *in vivo*. (A) Sample *D(t)* traces (top) and raster plots showing reconstructed CaT onsets (bottom). Note the developmental transition from discontinuous to continuous network activity. (B) Empirical probability distribution of CaT frequencies. (C) Mean CaT frequencies per FOV. (D) Lorenz curves of CaT frequencies. Line of equality (dotted) represents the case that all neurons have equal CaT frequencies. Inset depicts Gini coefficient calculation. (E) Mean Gini coefficients per FOV. (F) Cumulative probability of mean CV2 of inter-CaT intervals. Note that, at P11, CV2 distribution is narrower and centered around 1. (G) Mean CV2 per FOV. For a Poisson process, CV2 = 1 (dotted line). Each open circle represents a single FOV. Data are presented as mean ± SEM. ns – not significant. P4: P3–4, n = 19 FOVs from six mice, P11: P10–12, n = 11 FOVs from six mice, P18: P17–19, n = 12 FOVs from six mice, ^***^ P < 0.001. See also Table S2 and Source Data File.

Collectively, our data reveal a transient equalization in firing statistics of individual CA1 PCs during the second postnatal week, while highly skewed CaT frequency distributions eventually emerge only around/after eye opening.

### CA1 undergoes a transient enhanced bursting period while progressively transitioning from discontinuous to continuous activity

Previous *in vitro* work has identified giant depolarizing potentials (GDPs) as the most prominent pattern of correlated network activity in the neonatal hippocampus (Ben-Ari et al., 1989; Garaschuk et al., 1998; Leinekugel et al., 1997). GDPs depend on a depolarizing action of GABA_A_ receptor-dependent transmission (Ben-Ari et al., 1989; Owens et al., 1996) and disappear at around the beginning of the second postnatal week, when GABA actions shift from mainly excitatory to mainly inhibitory (Tyzio et al., 2007; Yamada et al., 2004). To investigate whether a similar developmental profile of NB generation exists in the CA1 *in vivo*, we next determined the time-course of the fraction of active cells Φ(t) (Fig. 3A). We found that, at P4, network activity was entirely discontinuous, as CA1 PCs spent relatively long time periods in a low-activity (silent) state, which was only interspersed by brief NBs (Fig. 3A, left, and Fig. 3B). During NBs, Φ(t) rarely exceeded 20% (Fig. 3B) indicating that the degree of co-activation *in vivo* is considerably lower than that reported for GDPs *in vitro* (Flossmann et al., 2019; Garaschuk et al., 1998; Leinekugel et al., 1997). In contrast to GDPs, bursting activity *in vivo* was even more pronounced at P11, when the network tended to oscillate between a silent state and a bursting mode with an inter-burst period of ∼2–10 seconds (Fig. 3A, middle). NBs were mainly generated during periods of discontinuous activity, which often lasted for several minutes. In addition, for the first time during development, continuous activity occurred, which we defined as prolonged non-zero activity (Figs. 3A and 3C and Methods). While CA1 started to dynamically transition between discontinuous and continuous activity at P11, the proportion of continuous activity substantially increased towards P18 (Fig. 3C).

**Figure 3.**
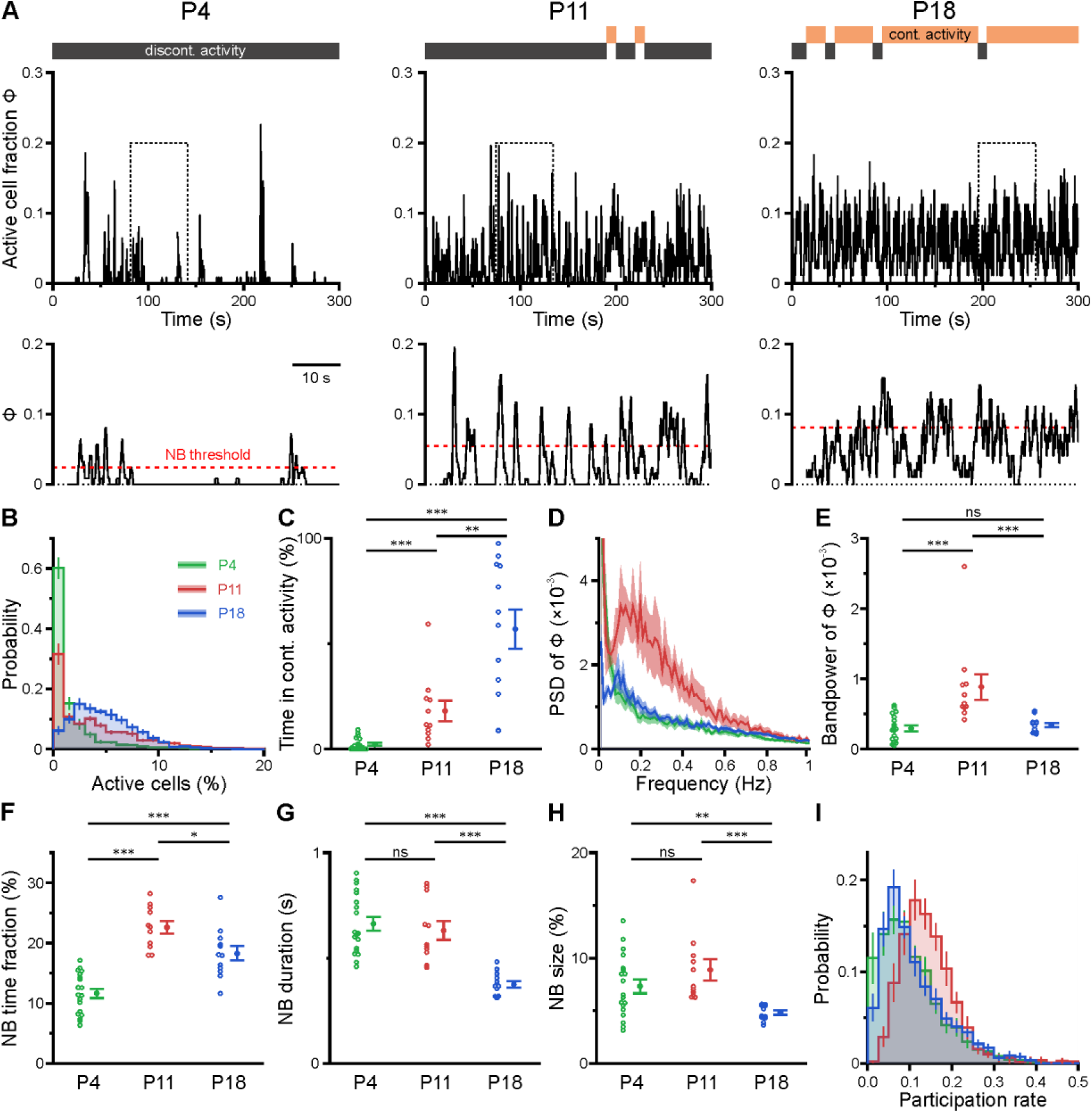
CA1 undergoes a transient enhanced bursting period while progressively transitioning from discontinuous to continuous activity. (A) Sample traces of the fraction of active cells Φ(t). Grey and orange bars indicate discontinuous and continuous network activity, respectively. Bottom traces show time periods marked on top (dotted rectangle) at higher temporal resolution. Red dotted lines indicate the activity-dependent thresholds for NB detection. (B) Empirical probability distribution of active cells per frame. (C) Continuous activity emerges at P11 and dominates over discontinuous activity at P18. (D) Power spectral density of Φ(t). (E) Bandpower of Φ(t) in the 0.1–0.5 Hz range. (F) The fraction of time that the network spends in NBs peaks at P11. (G) The average NB duration is lowest at P18. (H) Quantification of NB size as the mean fraction of active neurons (corrected for burst threshold as indicated in A). (I) Empirical probability distribution of the fraction of NBs that each cell is participating in. Each open circle represents a single FOV. Data are presented as mean ± SEM. ns – not significant. P4: P3–4, P11: P10–12, P18: P17–19, ^***^ P < 0.001. ^**^ P < 0.01. ^*^ P < 0.05. See also Fig. S1, Table S3 and Source Data File.

To quantify developmental changes in the rhythmicity of network activity, we first computed the power spectrum of Φ(t). At P11, this revealed a distinct peak in the range of ∼0.1–0.5 Hz (Fig. 3D), pointing to the existence of a preferred oscillation frequency of CA1 PCs. Such a power peak was absent at P4 and reduced at P18. Accordingly, band-power in the 0.1–0.5 Hz frequency range was significantly higher in the second postnatal week than at earlier or later stages (Fig. 3E, see Table S3).

To characterize periods of neuronal co-activation in more detail, we next extracted NBs by thresholding Φ(t). NB thresholds were determined separately for each FOV using surrogate data, accounting for differences in CaT frequencies (Fig. 3A, bottom, and Methods). The fraction of time that the network spent in NBs was lowest at P4 and peaked at P11 (Fig. 3F). Moreover, NBs at P18 were significantly shorter in duration than during the first and second postnatal weeks (Fig. 3G). We quantified NB size as the fraction of active cells (corrected for the threshold applied to Φ(t)) and found that it only declined after P11 (Fig. 3H). At P11, each neuron participated in 14.4 ± 1.2% of all NBs, which significantly exceeded participation rates at P4 (10.3 ± 0.8%) and P18 (11.2 ± 0.4%) (see #5 in Table S3). Importantly, these developmental changes in NB characteristics were robust to a wide range of NB definitions (Fig. S1 and Methods). Additionally, distributions of participation rates were relatively narrow at P11 (Fig. 3I), pointing to a greater similarity of cells with respect to their contribution to NB generation as compared to earlier or later developmental stages (#6 in Table S3).

Taken together, our data reveal that CA1 undergoes a transient period of enhanced bursting activity that coincides with the developmental appearance of continuous activity states during the second postnatal week. At this age, network activity displays rhythmicity in the sub-Hz range – in spite of the close-to-random activation of individual PCs.

### Enhanced population coupling underlies network burstiness in the second postnatal week in vivo

The transient developmental increase in bursting propensity was unexpected, as (1) GDPs *in vitro* disappear soon after the first postnatal week (Ben-Ari et al., 1989; Garaschuk et al., 1998; Khazipov et al., 2004) and (2) previous *in vivo* data from neocortex revealed a reduction in correlated network activity during the same time period (Colonnese et al., 2010; Golshani et al., 2009; Rochefort et al., 2009; van der Bourg et al., 2017). We therefore assessed whether the enhanced burstiness at P11 reflects an increase in functional neuronal coupling. We first investigated the coupling of single cell activity to that of the overall population. For each cell, we computed its population coupling (PopC) index (Okun et al., 2015; Sweeney et al., 2020) and tested for its significance using surrogate data (see Methods). The PopC index significantly peaked at P11 (Fig. 4A, Table S4), while there was no difference between P4 and P18. The higher PopC index at P11 arose from a significantly higher fraction of coupled cells (Fig. 4B), whereas the indices of significantly coupled cells were similar (Fig. 4C). We next addressed whether the increased PopC index at P11 results from an increase in pairwise temporal correlation of CaTs. To this end, we computed the spike time tiling coefficient (STTC) as a frequency-independent affinity metric of two event time series (Cutts et al., 2014) (Fig. 4D). The fraction of significantly correlated cell pairs did not significantly differ between P4 and P11 (P4: 13 ± 2%, P11: 23 ± 5%), but strongly decreased to 5 ± 1% at P18 (Fig. 4E). STTCs of significantly correlated pairs profoundly declined from 0.18 ± 0.01 at P4 to 0.09 ± 0.01 already at P11, but did not significantly change afterwards (P18: 0.10 ± 0.00; Figs. 4F and 4G). These data suggest that developmental changes in pairwise neuronal correlations do not account for the increased PopC nor the increased burstiness of CA1 PCs during the second postnatal week. We further analyzed the spatial structure of CA1 ensemble dynamics. We found that the dependence of STTCs on the Euclidean somatic distance was weak already during the first two postnatal weeks and non-significant at P18 (Fig. 4H), indicating that the horizontal confinement of patterned network activity is weak or absent throughout the developmental period studied here.

**Figure 4.**
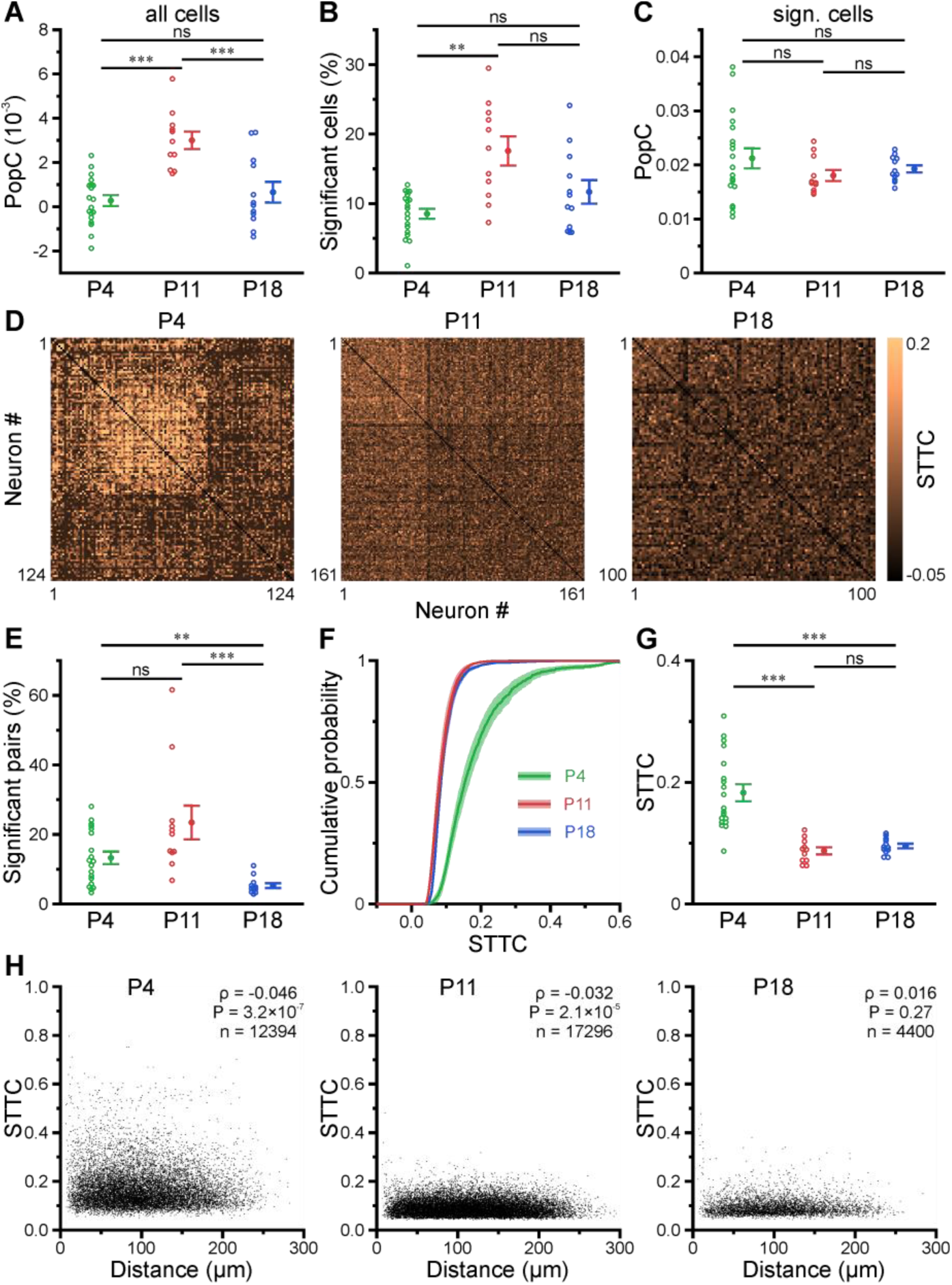
Enhanced population coupling underlies network burstiness in the second postnatal week *in vivo*. (A) The mean population coupling (PopC) index peaked at P11. (B) Mean fraction of cells with significant PopC. (C) Mean PopC index of significantly coupled cells only. (D) Sample STTC matrices (re-ordered). (E) Mean fraction of cell pairs having a significant STTC. (F) Cumulative probability of STTCs of significantly correlated cell pairs only. (G) Mean STTCs of significantly correlated cell pairs. (H) Relationship between STTC and Euclidean somatic distance for significantly correlated cell pairs. ρ denotes the Spearman’s rank correlation coefficient for all cell pairs analyzed (n) at a given age. (A–C and E–G) Each open circle represents a single FOV. Data are presented as mean ± SEM. ns – not significant. P4: P3–4, P11: P10–12, P18: P17–19, ^***^ P < 0.001. ^**^ P < 0.01. See also Table S4 and Source Data File.

Collectively, our data reveal that enhanced network burstiness during the second postnatal week is associated with a higher fraction of cells significantly locked to the activity of the local network, while pairwise neuronal correlations are low.

### Motifs of CA1 network activity undergo distinct developmental alterations

Recurring spatiotemporal cellular activation patterns are a hallmark of network activity in the adult hippocampus *in vivo* (Villette et al., 2015). Whether such repeating patterns (hereafter referred to as ‘motifs’) are already present at early developmental stages is unknown. To detect motifs, we divided the recording time into non-overlapping bins, each represented by a binary spatial pattern (vector) of active and inactive cells, followed by computing the matching index matrix of all possible pattern pairs (Fig. 5A). We then applied an eigendecomposition-based clustering method to each similarity matrix in order to detect potential motifs, while testing for their significance using surrogate data (see Methods). First, this analysis revealed that the global similarity of the activation patterns was lowest at P11 (Fig. 5B, Table S5), whereas it was similar between P4 and P18. This finding implies that there is less commonality between the sets of active cells present in different patterns at P11, and thus a more random recruitment of cells. Furthermore, the number of motifs was significantly lower at P11 (2.5 ± 0.9) as compared to P4 (5.9 ± 0.4) and P18 (7.0 ± 1.1) (Fig. 5C). When computing the fraction of patterns belonging to each motif, we found that the motifs had the highest repetition rate at P18 (32.0 ± 4.7%), while there was no significant difference between P4 (17.8 ± 1.2%) and P11 (16.6 ± 5.9%) (Fig. 5D). Taken together, these results suggest that recurring cellular activation patterns become more stable only after the onset of environmental exploration.

**Figure 5.**
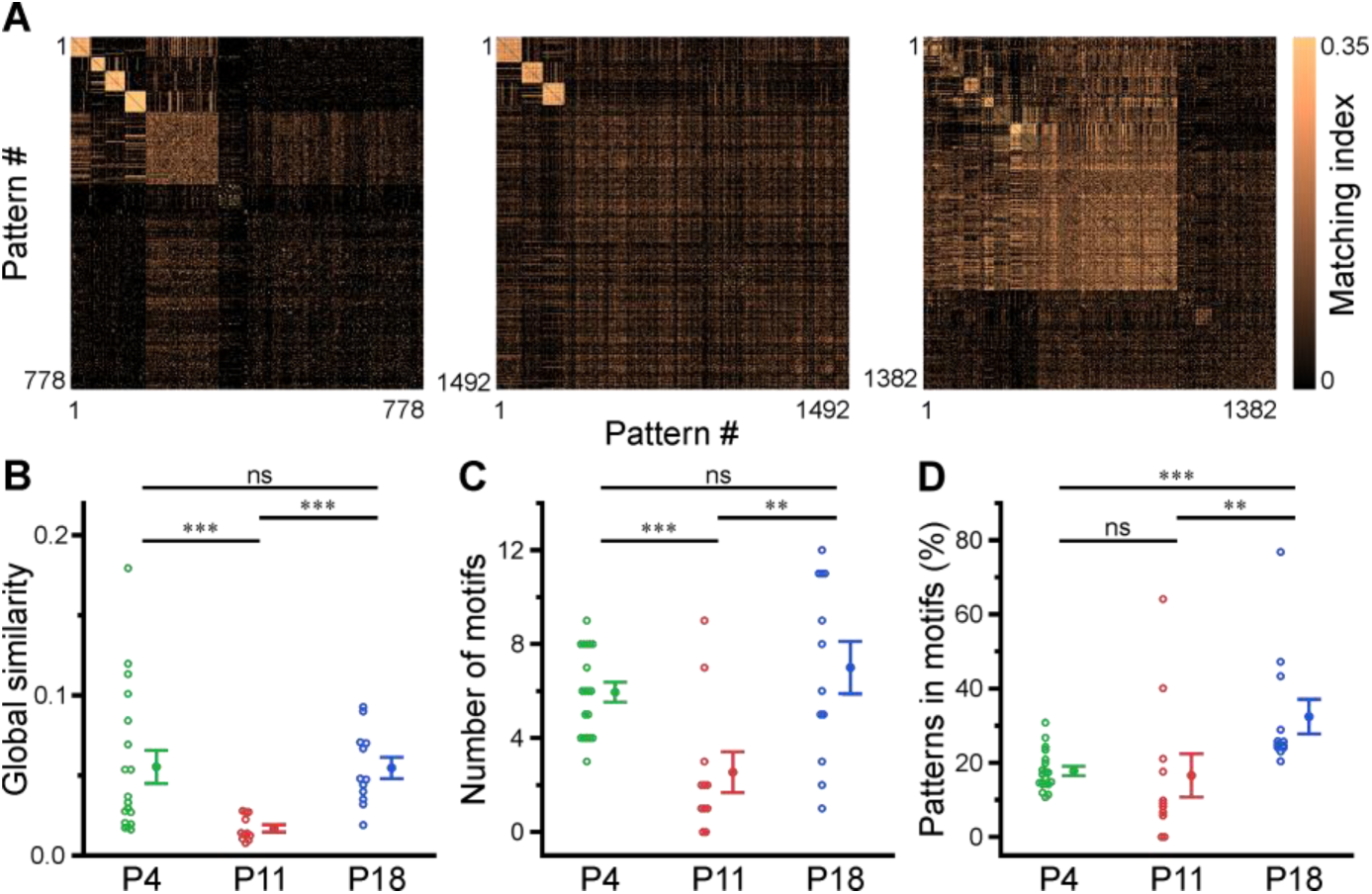
Motifs of CA1 network activity undergo distinct developmental alterations. (A) Similarity matrices (matching index) of binary activity patterns (re-ordered for illustration of motif detection). (B) Global similarity of activity patterns is lowest at P11. (C) The absolute number of detected motifs per FOV is lowest at P11. (D) Motif repetition quantified as the fraction of activity patterns belonging to each motif. Each open circle represents a single FOV. Data are presented as mean ± SEM. ns – not significant. P4: P3–4, P11: P10–12, P18: P17–19, ^***^ P < 0.001. ^**^ P < 0.01. See also Table S5 and Source Data File.

### Effects of nitrous oxide on body movements, vital parameters and network dynamics

In an attempt to reduce spontaneous movements and thus to increase mechanical stability during two-photon imaging, all measurements discussed so far were performed in the presence of the analgesic-sedative nitrous oxide, following our established procedures in neonatal mice (Kirmse et al., 2015). However, general anesthetics can have a profound impact on brain activity in age- and dose-dependent manner (Chini et al., 2019; Yang et al., 2021). As effects of nitrous oxide on neuronal dynamics in the developing CA1 had been unknown, we performed a separate set of experiments (n = 12 FOVs from six mice) in which we compared nitrous oxide to unanesthetized conditions using a paired design at P11 (Fig. S3A). Nitrous oxide reduced the time that mice spent in locomotion periods by several-fold (Figs. 6A, 6D and S2) and, hence, minimized periods of z-drift that needed to be discarded from analysis in two-photon Ca^2+^ imaging. Unlike most conventional anesthetics, however, nitrous oxide did not affect respiration or heart rate (Figs. 6B, 6C and S2). We observed slightly higher mean CaT frequencies in unanesthetized mice (Figs. 6E, S3B and S3C), while the oscillatory dynamics of CaTs (Figs. 6F and S3D) and NB properties (Figs. 6G and S3E) were unaltered. The most consistent effect of nitrous oxide was an increase in the percentage of correlated neuron pairs (based on STTC; Fig. 6H). This further translated into a higher fraction of cells that are significantly locked to local CA1 activity under nitrous oxide as compared to unanesthetized conditions (Fig. S3G).

**Figure 6.**
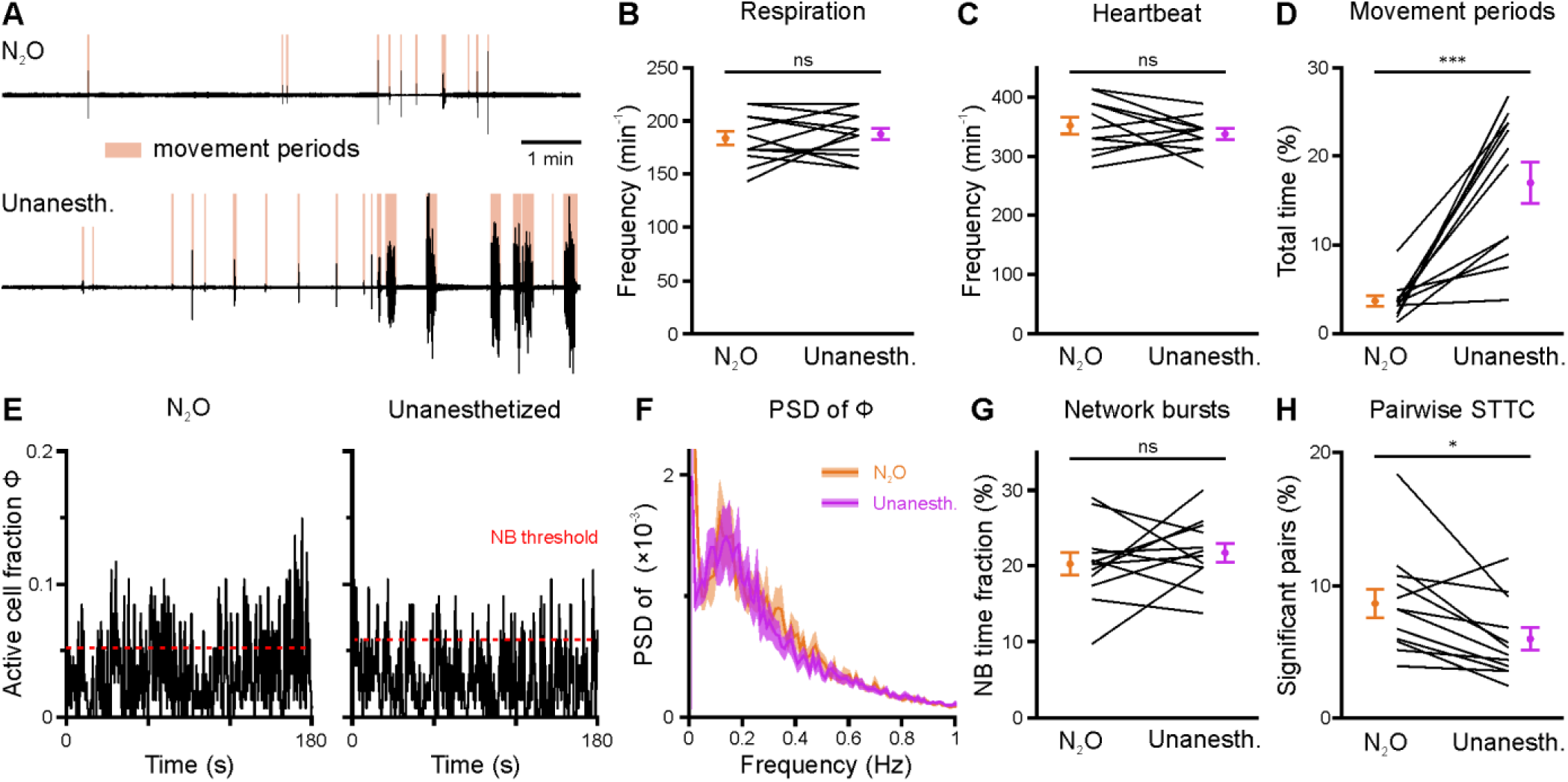
Effects of nitrous oxide (N_2_O) on body movements, vital parameters and network activity at P11. (A) Sample respiration/movement signal from an individual mouse receiving either 75% N_2_O/25% O_2_ (*top*) or pure O_2_ (Unanesth., *bottom*). Detected movement periods are highlighted. (B) Respiration rate is unaffected by N_2_O (n = 12 FOVs from six animals). (C) Heart rate is unaffected by N_2_O (n = 11 FOVs; in the recording from one FOV, heart rate could not be reliably determined). (D) N_2_O significantly reduces movement periods. (E) Sample traces of the fraction of active cells Φ(t) (Δt = 3 frames). Red dotted lines indicate the activity-dependent thresholds for NB detection. (F) Power spectral density of Φ(t) is similar in the presence or absence of N_2_O. (G) Fraction of neuron pairs having a significant STTC. (H) The fraction of time that the network spends in NBs. n = 12 FOVs from six mice. ^***^ P < 0.001, ^**^ P < 0.01, ns – not significant. See also Table S6 and Source Data File.

In sum, neuronal network activity under nitrous oxide closely resembles that recorded in unanesthetized mice despite minor quantitative differences. A major advantage of its use lies in the reduction of extended movement periods, which significantly increases mechanical stability during two-photon imaging.

### A neural network model with inhibitory GABA identifies intrinsic instability dynamics as a key to the emergence of network bursts

Hitherto our analyses of experimental data revealed an unexpected bursting behavior of CA1 PCs at P11, despite the developmental emergence of synaptic inhibition (Murata et al., 2020; Spoljaric et al., 2017; Tyzio et al., 2007), which we found to associate with their higher coupling to local network activity. However, the mechanisms governing *in vivo* network burstiness as well as its functional implications remain to be understood. Here, we provide mechanistic insights into these open questions by using computational network modeling and stability analysis techniques.

We employed a recurrent neural network (RNN) model of mean firing-activity rates of excitatory glutamatergic (PC) and inhibitory GABAergic (IN) cell populations (*A*_PC_ and *A*_IN_) with dynamic synaptic weights (Flossmann et al., 2019; Rahmati et al., 2017) (Fig. 7A). Here, we constrain the model with previously reported and our present experimental data obtained for P11: I) GABAergic synapses are considered to be inhibitory (Kirmse et al., 2015; Murata et al., 2020; Valeeva et al., 2016) and II) the spontaneous time-averaged *A*_PC_ is effectively non-zero (Fig. 2C). We found that such a network operates under a bi-stable regime, where two stable spontaneous fixed points (FPs) exist: one at a silent state (*A*_PC_ = *A*_IN_ = 0 Hz) and the other at an active state (*A*_PC_ ≠ 0 Hz and *A*_IN_ ≠ 0 Hz; green dots in Fig. 7B). This is reminiscent of our experimental observations demonstrating that CA1 dynamically transitions between discontinuous and continuous activity states at this stage (see Figs. 3A–C). The ability of the network model to embed the latter FP is mainly due to the stabilization function of inhibitory GABA (Latham et al., 2004; Rahmati et al., 2017). Strikingly, our simulations showed that the network can process a given input quite differently at the silent and active states, respectively (time points a and c in Fig. 7C). To this end, we applied a set of two excitatory inputs to the network’s PC and IN populations (*e*_PC_ and *e*_IN_), resembling e.g. SPW-driven inputs to CA1 (Fig. 7C). We set the input strengths and duration to be identical across the two states. We found that, when operating at the active state, the network activity monotonically decays back to this state, once the input ceases (a in Fig. 7C). However, at the silent state, input removal is followed by a transient profound surge in network activity (c in Fig. 7C). Hereafter, we refer to this supra-amplification activity as simulated NB (simNB), emulating experimentally observed NBs (Fig. 3).

**Figure 7.**
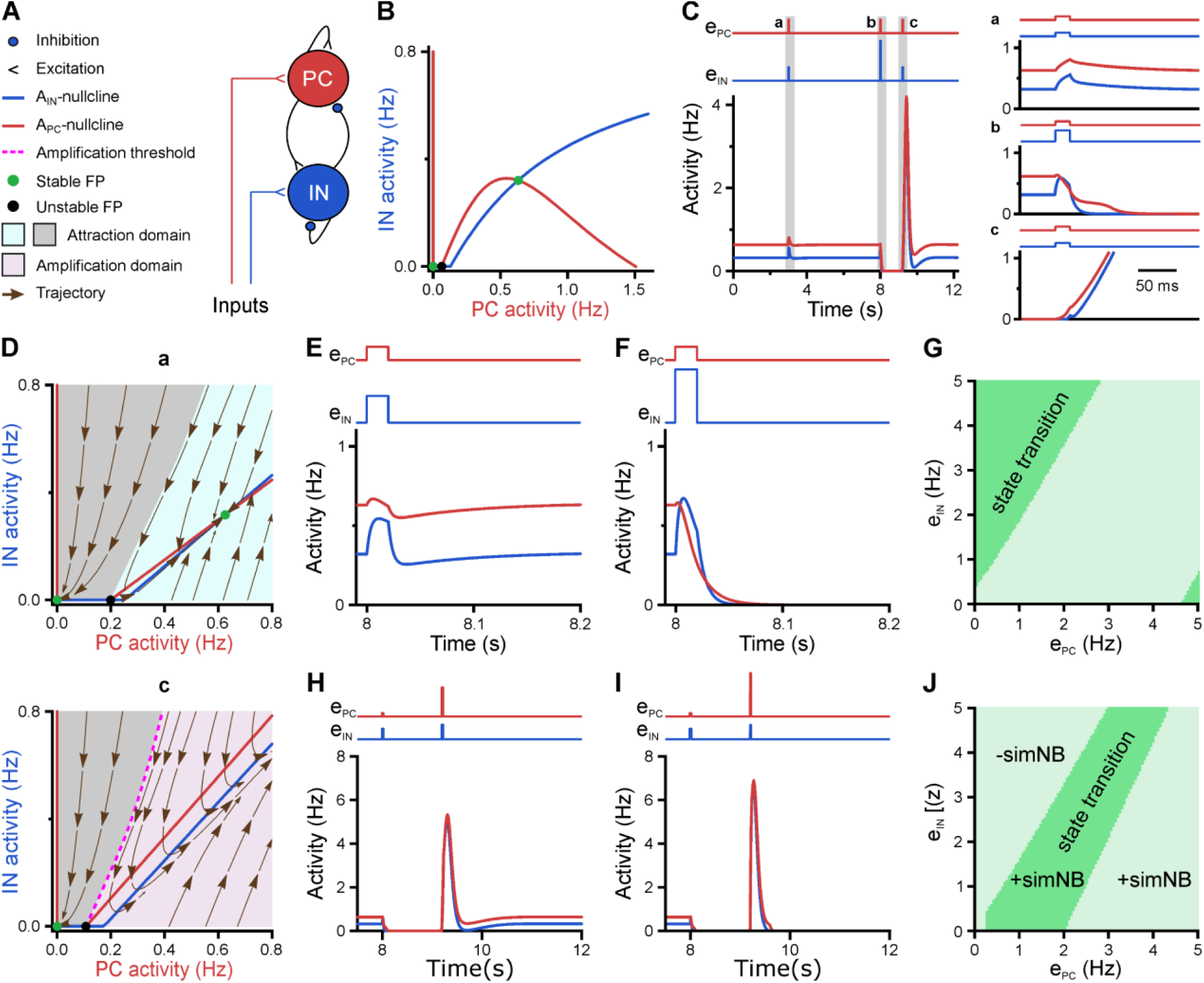
A neural network model with inhibitory GABA identifies intrinsic instability dynamics as key to the emergence of network bursts. (A) Schematic diagram of the STP-RNN model. (B) The *A*_IN_ -*A*_PC_-plane of the full STP-RNN’s stationary dynamics. Note the presence of two stable fixed points (FPs; green dots) at silent and active states as well as the unstable FP (black dot). (C) simNB generation requires network silencing. The model was stimulated by pulse-like input to both PC and IN populations for a duration of 0.020 s (at t = 3 and 9.2 s: *e*_PC_ = *e*_IN_ = 0.25; at t = 8 s: *e*_PC_ = 0.25, *e*_IN_ = 0.75). Zoom-in of the activity around the stimulation times at active (a and b) and silent (c) states are shown in right panels. Input time series are shown on top of the plots. (D) The presence of an amplification domain in the initial phase of network firing dynamics enables the emergence of simNBs. The *A*_IN_ -*A*_PC_-plane of the STP-RNN with synaptic efficacies frozen at active (a, top) and silent (b, bottom) states, right before input arrival. (E–G) Transition from active to silent state requires specific input ratios. Input delivered at t = 8 s. (E) A failed transition: *e*_PC_ = 0.25, *e*_IN_ = 0.5. (F) A successful transition: *e*_PC_ = 0.25, *e*_IN_ = 1. (G) A color-coded matrix of successful (dark green) and failed (light green) transitions to the silent state in response to different combinations of *e*_PC_ and *e*_IN_ amplitudes. (H–J) Both the transition from the silent to the active state and the simNB generation require specific input ratios. Input delivered at t = 9.2 s. Same as E–G, but for the backward transition to the active state. +simNB and – simNB indicate the emergence and absence of bursts.

### Network state-dependency of simNB generation

To disclose the mechanisms underlying this distinct behavior of the network at the silent and active state (Fig. 7C), we computed the corresponding steady-state *A*_IN_ -*A*_PC_ –plane of the network, after freezing the slow short-term synaptic plasticity (STP) dynamics and, thus, synaptic weights (Frozen STP-RNNs), at either of these states separately (Fig. 7D). This analysis enables assessing the initial phase of network activity following an input perturbation. We found that, while operating at the silent state, the active state is not initially accessible to the network (lower panel in Fig. 7D). Instead, an unstable FP is present in the network’s fast (i.e. firing activity) dynamics, which builds an amplification threshold around the attraction domain of the FP located at the silent state. This in turn allows for the emergence of simNBs: A sufficiently strong perturbation, amenable to initially push the network activity beyond this threshold (i.e. to the amplification domain), will transiently expose the network to its intrinsic instability-driven dynamics, thereby effectively triggering a simNB (c in Fig. 7C). Note that this unstable FP is different from its counterpart in the full system (Fig. 7B) and is only visible in the network’s fast dynamics. In particular, this FP is transient and disappears around the peak of the elicited simNB, mainly due to short-term synaptic depression (Rahmati et al., 2017). Unlike the silent state, the network frozen at the active state has no amplification domain, but instead two attraction domains related to its FPs at silent and active states (upper panel in Fig. 7D). This explains the network’s incapability of eliciting simNBs when operating at the active state. Collectively, these results suggest that simNBs, initiated by the input, are mainly an expression of the network’s intrinsic instability dynamics, where the silent periods of the network are a prerequisite for its emergence. The model behavior is in agreement with our experimental data indicating that NBs are chiefly generated during periods of discontinuous activity (see Figs. 3A– C).

### Input-strength dependency and internal deadline of state transitions

What are the input requirements that allow the network to transition between the active and the silent states? First, we found that the silencing of the network in an active state requires specific ratios of excitatory input strengths to be delivered to its PC and IN populations (Figs. 7E–G). In particular, the presence of GABAergic inhibition can effectively promote this transition, where otherwise a relatively much stronger *e*_PC_ is required to silence the network solely (Fig. 7G). Furthermore, once silenced, pushing the network back to the active state is also dependent on input ratio (Figs. 7H–J). However, to make such a transition, the network becomes noticeably more selective about the input ratio (compare Figs. 7G and 7J). For both transitions, the proper ratios of the inputs are effectively determined by the approximated initial phase of the network response (Fig. 7D), and thus mainly dependent on the synaptic weights right before the input arrival. In sum, these results suggest that proper input strengths onto the PC and IN populations, along with the inhibitory action of GABA, play key roles in the dynamic state transitioning of the network, thereby allowing for its burstiness.

Considering the dynamics of synaptic weights in our model along with their significance for state transitions, we next investigated the impact of input timing (Fig. 8). We found that, once silenced by the first input, a deadline is formed for the network’s transitioning back to the active state (dotted line in Figs. 8A, 8D, 8G and 8H). If the second input misses the deadline, the network will elicit a large-amplitude simNB, but is not able to converge to the active state any longer (Fig. 8D). Prior to this deadline and depending on the input ratio (Fig. 7J), the network will either transition to the active state (Figs. 8A and 7H) or return to the silent state (Fig. 7I). Importantly, our analysis showed that this deadline is an internal property of the network and cannot be overruled by any input level (see below). Therefore, specific combinations of input ratio (Fig. 7J) and input timing (Fig. 8G) are required for transitioning to the active state. In addition, once the simNB failed to converge to the active state, the network will encounter a new deadline (Fig. S4). In sum, these results imply that the silent state of the network can have *per se* different hidden sub-states, each with a specific input-encoding operating scheme.

**Figure 8.**
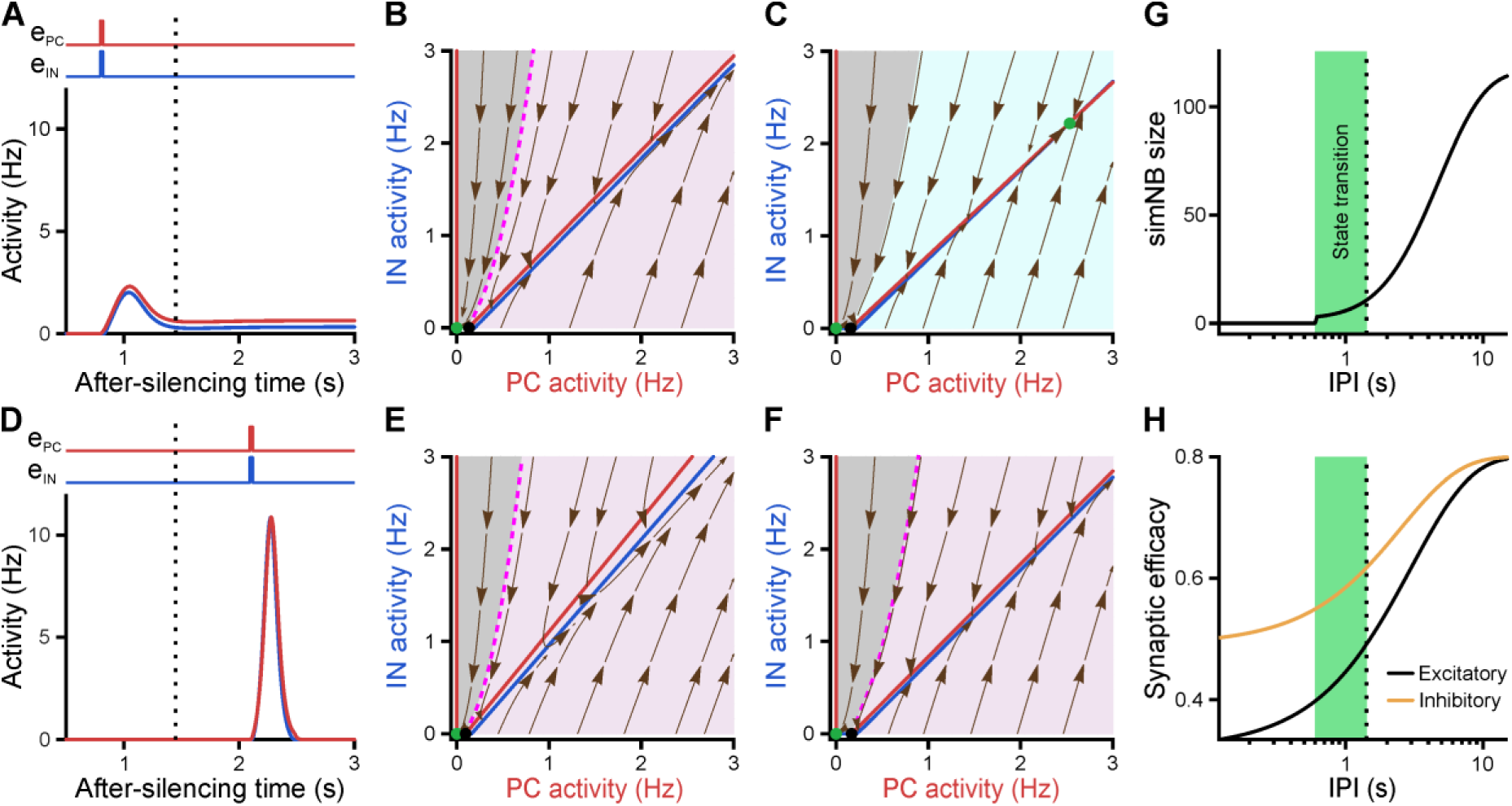
Internal deadline of state transitions. (A–C) Input delivered to the network before the deadline can move it to active state. (A) A successful transition. The input delivered at t = 0.8 s; *e*_PC_ = 0.25, *e*_IN_ = 0.25. (B) The *A*_IN_ -*A*_PC_ –plane of the STP-RNN with synaptic efficacies frozen at the silent state right before the input arrival. (C) Same as B, but frozen at the peak of the network burst (i.e. simNB) shown in A. Note the presence of the transient stable FP (non-origin green dot), which triggers the transitioning to the active state. (D–F) Once the deadline is missed, the network cannot be moved to the active state by the subsequent input. Same as A– C, but the input delivered at t = 2.1 s. Note the absence of a non-origin transient stable FP in F, in contrast to C. (G) The simNB size and network transition to the active state depend on the inter-pulse intervals (IPI: the arrival time of the next input relative to the silencing time of the network). simNB size is computed as the maximum of *A*_IN_ + *A*_PC_ after the secondary input. Note the presence of a short window for transitioning to the active state. *e*_PC_ = 0.25, *e*_IN_ = 0.25. (H) Same as G, but for the non-scaled efficacies of GABAergic (*u_I_ x_I_*; orange; see Methods) and glutamatergic (*u_p_ x_P_*; black) synapses, right before the arrival of the secondary input. (A, D, G, H) The dotted line at t = 1.45 s depicts the internal deadline.

Having found the intrinsic deadline as a main determinant for the type of network burst, we next investigated the origin of these different activity patterns: How does the network decide between transitioning to the active state and returning to the silent state? Remarkably, we found that the deadline for network transitioning to the active state is mechanistically dependent on the presence of a transient stable FP in its fast dynamics around the peak of the simNB. This can be seen in the two examples where the network receives the same input but at different inter-pulse intervals (IPIs), one preceding (Figs. 8A–C) and the other exceeding the deadline (Figs. 8D–F). For both IPIs, at the time right before the second input (Figs. 8B and 8E), the Frozen RNNs only provide evidence for the emergence of simNB, but not for the state transition (note the presence of an amplification domain; pink area). Importantly, we found, however, that in the case of the shorter IPI, the network is able to form a transient, stable non-zero FP in its Frozen RNN, at the peak of the simNB (compare Figs. 8C and 8F). This FP can transiently attract the network’s activity towards itself, and as the activity evolves accordingly, it also changes its position in the corresponding updated Frozen STP-RNN, until eventually converging to its counterpart in the full system. Intuitively, this transient, stable FP can guide the network’s activity towards that of the full system (see the non-origin green dot in Fig. 7B). The temporal repositioning of this stable FP is due to the activity-dependency of the synaptic weights in our model. Besides, our findings show that the existence of this FP around the simNB peak is effectively determined by simNB size (Fig. 8G). If simNB size exceeds an internally determined threshold, the network cannot build such a transient stable FP due to a reduction of synaptic weights (Rahmati et al., 2017); consequently, the simNB will be attracted towards the silent state. In this line, Fig. 8G shows that simNB size is effectively determined by the IPI: The longer the IPI (thus, the silent period) is, the larger the simNB will be. Here, the IPI-dependency of the simNB size mainly reflects the slow recovery from short-term depression of excitatory synapses at the silent state (Fig. 8H).

In conclusion, our modeling results indicate that developing CA1 possesses multiple input-encoding schemes, which are effectively determined by three factors: 1) the input ratio, 2) the input timing, and 3) the non-linearity and dynamics of synaptic weights.

### A bi-stable STP-RNN model with inhibitory GABA robustly explains our experimental observations

We next investigated whether alternative network models (or mechanisms) are better suited to explain the observed CA1 dynamics in the second postnatal week. By decreasing the population-activity thresholds (θ) and/or changing the polarity of GABAergic synapses from inhibitory to excitatory, we created two operationally distinct models called Mono-RNNi (θ_PC_ ↓, ‘i’ for inhibitory GABA) and Mono-RNNe (J_IN_ → −0.5×J_IN_, θ_PC_ ↓, θ_IN_ ↓; ‘e’ for excitatory GABA) (Fig. S5A). Of note, lower θ reflects a higher background input to, or a lower spike threshold of, the neuronal population (see Methods). Each of the models is mono-stable with one spontaneous FP in an active state (non-origin green dots in Fig. S5A).

As a prerequisite for comparison, both Mono-RNNi and Mono-RNNe can generate silent periods, simNBs, and an active state with *A*_PC_ > 0 (Figs. S5B–F), as observed in our data. As compared to the bi-stable STP-RNN model (Figs. 7 and S5A), however, both mono-stable networks are less plausible in explaining our experimental observations for several reasons. First, due to the lack of a silent FP, Mono-RNNi and Mono-RNNe can transition to and remain in a silent state only in the (continuous) presence of synaptic input (Figs. S5C and S5D). Second, for silencing (*A*_PC_ =*A*_IN_ =0), the external input to at least one of the populations needs to be inhibitory (Fig. S5E, green areas). However, input to CA1 at this age is mainly mediated by glutamatergic projections from entorhinal cortex and CA3. In addition, considering inhibitory input violates the assumption of excitatory GABAergic synapses in the Mono-RNNe. Third, both Mono-RNNi and Mono-RNNe require a relatively long silencing (of at least the PC population) to effectively generate simNBs (Fig. S5F), which is difficult to reconcile with the time-course of SPWs (tens of milliseconds).

The intrinsic bi-stability of the STP-RNN renders it computationally different from the mono-stable alternatives: (i) In the STP-RNN, simNBs are triggered by suitable inputs (Figs. 7 and 8), whereas in Mono-RNNi and Mono-RNNe, simNBs are elicited in the form of rebound bursts only after the cessation of the non-specific silencing inputs (Figs. S5C, S5D and S5F). This feature renders the STP-RNN potentially more suitable for input discrimination prior to the onset of environmental exploration. (ii) Regardless of their size, rebound simNBs will always return to the active state in Mono-RNNi and Mono-RNNe (Figs. S5C and S5D), in contrast to the STP-RNN, which may also return to its rest state (Figs. 7 and 8). (iii) Mono-RNNe lacks an inhibition-stabilized network (ISN) regime (Fig. S5A), which is thought to provide a general strategy for supporting more complex computations (Latham et al., 2004; Tsodyks et al., 1997).

In sum, these results indicate that the proposed STP-RNN, operating under bi-stability, is not only computationally more flexible, but also more plausible in explaining our experimental observations.

### Inhibitory stabilization of a persistent active state in the bi-stable STP-RNN model

Using our STP-RNN model, we further investigated the functional significance of the network behavior during the second postnatal week. Our analyses show that an ISN regime is effectively accessible at this age, due to the developmental increase of synaptic inhibition (Figs. 9A and 9B). This presumably enables CA1 to process more complex computations (while avoiding instabilities) in parallel to the developmental strengthening of sensory inputs. Importantly, the active state in our model is also located in the ISN FP-domain (Fig. S5A, left). Such an ISN regime is absent if GABAergic transmission is excitatory (Figs. 9A (right) and 9B). Moreover, the developmental increase in the strength of synaptic inhibition (J_IN_) enables the network to operate as an ISN at a wider range of PC activity levels (Fig. 9C). This is because the unstable FP-domain, confined to low levels of PC activity, is progressively replaced by the ISN FP-domain as inhibitory synapses become stronger (Figs. 9A and 9B). In the model, simNB size is reduced by a strengthening of inhibition (akin to our experimental observations at P11 vs. P18), but increased if GABA is considered excitatory (unlike our data at P11 vs. P4) (Fig. 9D). This implies that the changes in inhibitory strength alone are unsuited to explain the enhanced burstiness in the second postnatal week. We found, however, that the experimentally reported, concurrent developmental strengthening of glutamatergic synapses (J_PC_) (for review see Kirmse et al., 2022) exerts an opposite effect by profoundly amplifying simNBs (Fig. 9E). Collectively, our analyses portend that the emergence of a persistent active state in CA1 reflects the developmental strengthening of both GABAergic inhibition (J_IN_) and glutamatergic excitation (J_PC_) as well as changes in background input and/or intrinsic excitability (θ). In line with this, the enhanced burstiness at P11 is an expression of the complex neural dynamics in a bi-stable STP-RNN that identify the second postnatal week as a key transitional period in CA1 network maturation.

**Figure 9.**
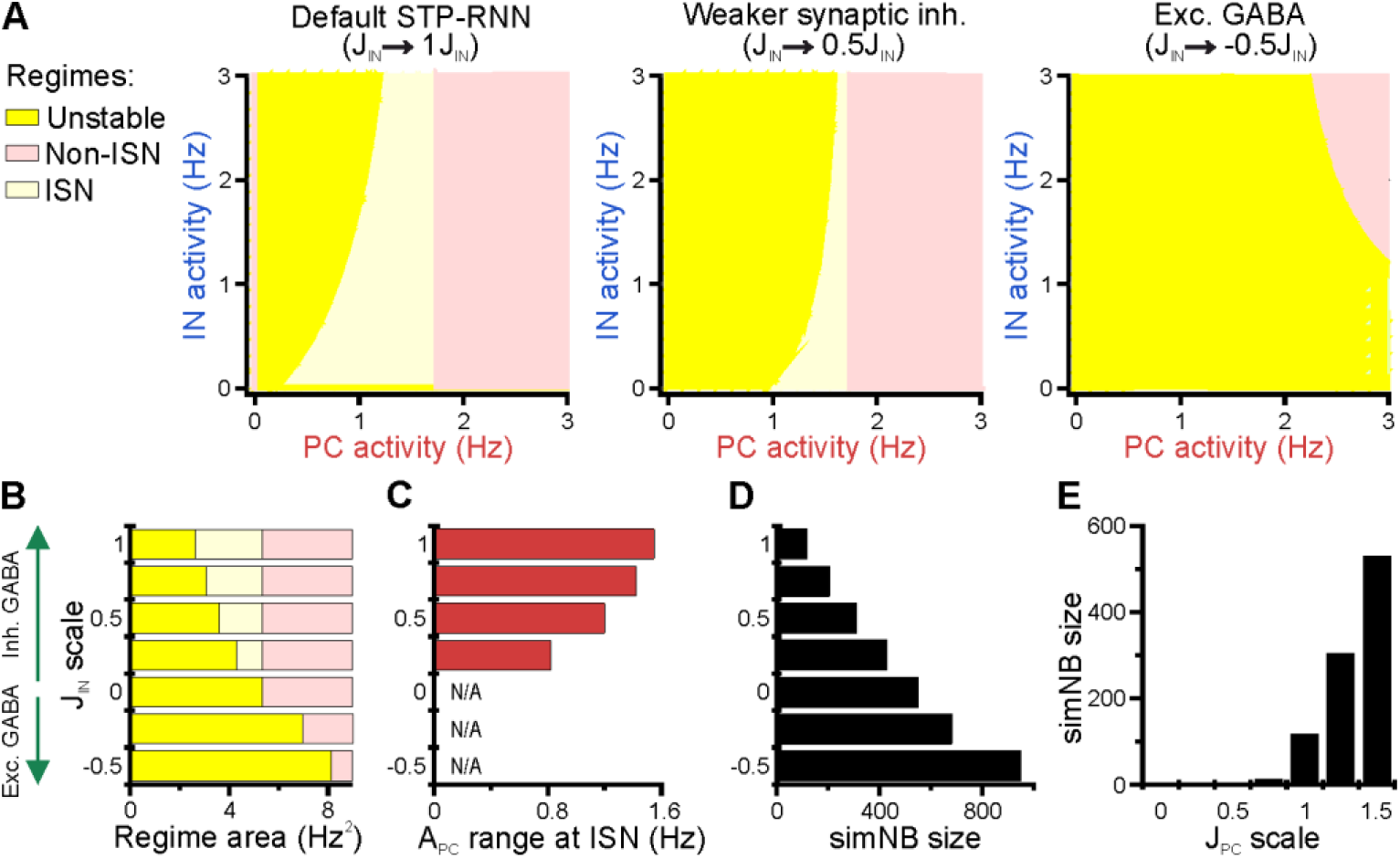
Inhibitory stabilization of a persistent active state in the bi-stable STP-RNN model. (A) The ISN regime becomes accessible to the network upon the developmental emergence of synaptic inhibition. The colored regions in each *A*_IN_ -*A*_PC_ –plane of the network model depict the FP-domains of three possible operating regimes: unstable dynamics, ISN, and Non-ISN. Synaptic inhibition strength J_IN_ was set to 3 (inhibitory; *left*), 1.5 (inhibitory, *middle*), and −1.5 (excitatory, *right*). (B–D) Effect of strength and polarity of GABAergic synapses (J_IN_) on the availability of the ISN regime (B), the maximum range of PC activity in the ISN regime (C), and size of simNBs when triggered at the rest state (D). Results were obtained by scaling J_IN_ in the STP-RNN model with values indicated on the y-axis. The area of each operating regime was computed based on the area of its FP-domain in the *A*_IN_ -*A*_PC_ –plane (with limits as in A). (E) Dependency of simNB size on the strength of glutamatergic synapses (J_PC_). Default value of J_PC_ in STP-RNN was 6.5 (corresponding to a scale of 1).

## Discussion

### Developmental trajectories of network dynamics in CA1 in vivo

Our data demonstrate that activity of CA1 PCs in the first postnatal week is generally low and exclusively discontinuous in nature, as they spend most of the time in a silent rest state that is only interrupted by relatively brief NBs (Figs. 2A–C and 3A–C). In other words, CA1 PCs are incapable of sustaining persistent activity at this age, similar to findings in visual cortex (Ackman et al., 2012; Hanganu et al., 2006; Kirmse et al., 2015; Kummer et al., 2016). Previous data-driven computational modeling further showed that such dynamics can be emulated by mono-stable network models, in which excitatory GABA effectively promotes network burstiness (Flossmann et al., 2019). CA1 NBs differ from their neocortical counterparts in that the latter exhibit a distinct horizontal confinement that appears to be largely absent in CA1. This is evident in the weak distance-dependency of pairwise correlations of CA1 PCs (Fig. 4H), extending previous results from large-scale imaging (Graf et al., 2021) and electrophysiological (Valeeva et al., 2019; Valeeva et al., 2020) studies. The developmental relevance of such differences in spatial characteristics is unknown. However, wavefront-containing activity patterns have been causally linked to the developmental refinement of topographic maps (Cang et al., 2005; Li et al., 2013) and receptive field characteristics (Albert et al., 2008; Wosniack et al., 2021) in the visual cortex, suggesting that their absence in CA1 might reflect the lack of a clear topical macro-organization of the hippocampus (Bellistri et al., 2013).

We here identify the second postnatal week as a key transitional period in CA1 network maturation. For the first time in development, CA1 PCs are able to maintain spontaneous persistent activity, while this transition towards embedding continuous activity is largely completed by P18 (Figs. 3A–C). In the second postnatal week, CA1 PCs undergo a transient period of enhanced network burstiness, in which NBs chiefly arise during periods of discontinuity (Fig. 3A). This trajectory remarkably differs from what has been previously reported for the hippocampus *in vitro*, where GDPs disappear shortly after the first postnatal week. At this time, GABA-releasing INs already inhibit CA1 PCs (Murata et al., 2020; Spoljaric et al., 2017; Tyzio et al., 2008), implying that NBs *in vivo* do not depend on GABAergic excitation (in contrast to GDPs). NBs are also observed at P18, but these are short in duration (Fig. 3G) and recruit fewer neurons (Fig. 3H) than at earlier stages. Strikingly, whereas NBs at P11 are large and frequent, recurring cellular activation patterns (a prime characteristic of adult hippocampal activity) become more stable only after the onset of environmental exploration (Fig. 5D). This was accompanied by the developmental emergence of highly skewed firing rate distributions by P18 (quantified as CaT frequencies; Figs. 2B–E). The latter are thought to underlie sparse coding (Ikegaya et al., 2013; Narayanan et al., 2012; Roxin et al., 2011; Trojanowski et al., 2020; Yassin et al., 2010), an energy-efficient regime of input processing and information storage (Mizuseki et al., 2013). Collectively, our data indicate that CA1 network activity acquires a number of “adult-like” characteristics by P18, i.e. shortly after the onsets of pattern vision, active whisking and environmental exploration.

### A role for intrinsic network instability and synaptic inhibition in NB generation in CA1

During the transition period (i.e. P11), bursting activity exhibited a preferred frequency of ∼0.1–0.5 Hz, indicating that NBs occur in a temporally non-random manner (Fig. 3). However, individual neurons were recruited more randomly at this age, as the number of significant motifs of network activity as well as their average repetition probability were lowest (Fig. 5). At P11, the occurrence of CaTs in individual cells resembled a Poissonian process (Fig. 2), and pairwise neuronal correlations were lower than at P4 (Fig. 4G). We here set out to explain these seemingly discordant experimental findings using data-informed computational modeling.

Capitalizing on a dynamic systems modeling approach, we show that a potential dynamical regime of the network that allows for the generation of NBs in the presence of effective synaptic inhibition is bi-stability. We found that our network model is prone to an intrinsic instability, governed by a nonlinear interaction between its fast (firing) and slow (synaptic) dynamics. Such instability enables the model to over-amplify the input, even after its removal, and thus elicit network bursts (simNBs; Fig. 7). This indicates that a (sim)NB reflects a spatiotemporal trajectory of the network’s intrinsic instability dynamics, which, due to its nature, can recruit a random set of cells at random order within a specific time-window. The size of this set and the time-window are determined by the synaptic weights right before the input arrival (Rahmati et al., 2017). Thus, the data-informed model mechanistically links strong population coupling to weak pairwise neuronal correlations, the close-to-random firing of individual PCs and the low number of network motifs – as we found experimentally for the second postnatal week.

What are the functional roles of burstiness and synaptic inhibition at this stage? Our model, in addition to its silent state, embeds a stable fixed point (or steady state) at non-zero low activity rates (Fig. 7B), in accordance with our recorded data. Theoretical studies indicate that the presence of such a fixed point requires the stabilization function of inhibitory GABA (Latham et al., 2004; Ozeki et al., 2009; Rahmati et al., 2017; Tsodyks et al., 1997). We here show that, at such a fixed point, the analyzed network model operates under an ISN regime, which may enable CA1 networks to begin performing complex computations (Latham et al., 2004; Tsodyks et al., 1997). We also show that the ISN FP-domain becomes effectively expanded by the strengthening of inhibition, i.e. a larger set of stimulus-evoked ISN attractors (or FPs) are accessible for network computations, in parallel to the developmental strengthening of sensory inputs. The ability of the network to dynamically transition between its silent and active states in an input-dependent fashion (Fig. 7) renders the second postnatal week an early developmental stage toward forming hippocampal memory and cognition mechanisms, as found in adult hippocampal attractor networks (Hartley et al., 2014; Knierim et al., 2016; see also Rahmati et al., 2017; Rolls, 2007). This view is supported by (I) the existence of the internal deadlines as well as a delicate input-ratio and -timing dependency of successful state transitions and simNB generation and (II) the network’s ability to store information in both the silent and active state through transient synaptic weights (Barak et al., 2014; Mongillo et al., 2008; Stokes, 2015) and persistent activity (Boran et al., 2019; Zylberberg et al., 2017). Our modeling results further imply that the network’s silent state has *per se* several dynamic operational sub-states, which keep track of input timing and strength (Figs. 8 and S4) to produce proper network read-outs. Collectively, we postulate that the basis of CA1 encoding schemes is set in shortly before eye opening. Moreover, our data suggest that GDPs disappear due to improper synaptic inputs *in vitro* during the second postnatal week, when GABA actions switch to inhibitory (Murata et al., 2020; Valeeva et al., 2016).

### Potential developmental functions of network bursts in the neonatal CA1

Computational modeling suggests a mechanism, whereby CA1 undergoes extensive input-discrimination learning before eye opening. In this scenario, NBs serve as a feedback that informs individual CA1 PCs about functionally important characteristics of the synaptic input to the local network, including (I) the proper targeting ratio of excitatory PCs *versus* inhibitory GABAergic INs (Murata et al., 2020; Valeeva et al., 2016) and (II) the timing of inputs relative to the network’s operational state. Interestingly, the developmental period of enhanced network burstiness coincides with a major surge of synaptogenesis in CA1 PCs (Kirov et al., 2004). The latter involves a net addition of synapses, but also functionally important anatomical rearrangements. Specifically, the formation of mature dendritic spines, which allow for electrical and metabolic compartmentalization of postsynaptic responses, commences only at around P10, by which time most glutamatergic synapses are rather localized to dendritic shafts (Fiala et al., 1998; Kirov et al., 2004). In addition to acting as potential synaptogenic stimuli (Kirov et al., 2004), NBs could thus be an important element underlying synaptic competition and pruning, for example, based on synchronization-dependent plasticity rules in nascent dendrites (Winnubst et al., 2015). Network burstiness might therefore be causally related to the delayed development of skewed (approximately log-normal) firing rate distributions (Fig. 2) underlying sparse coding (Ikegaya et al., 2013; Narayanan et al., 2012; Roxin et al., 2011; Trojanowski et al., 2020; Yassin et al., 2010). In accordance with the efficient coding hypothesis and seminal work in the visual system (Albert et al., 2008), we argue that one function of developing CA1 and, thus, NBs is to remove statistical redundancy in the multi-sensory place-field code, by making use of a learning scheme that uses both intrinsically and sensory-evoked activity already before environmental exploration.

## Acknowledgments

We thank Ina Ingrisch for excellent technical assistance. This work was supported by Individual Research Grants (KI 1816/6-1, KI 1816/7-1 to K.K., HO 2156/5-1, HO 2156/6-1 to K.H.), the Research Unit 3004 (KI 1816/5-1 to K.K., GE 2519/8-1, GE 2519/9-1 to C.G.), the Priority Program 1665 (HO 2156/3–1/2 to K.H., KI 1816/1–1/2 to K.K., KI 1638/3–1/2 to S.J.K.) and the CRC Transregio 166 (B2 to C.G., B3 to K.H., K.K.) of the German Research Foundation.

## Author Contributions

Conceptualization, K.K., K.H., V.R.; Methodology, J.G., K.K., V.R.; Formal Analysis, K.K., J.G., V.R.; Investigation, J.G., K.K., V.R., M.M.; Computational modeling, V.R.; Writing – Original Draft, K.K., V.R., J.G.; Writing – Review & Editing, J.G., V.R., M.M., O.W.W., C.G., S.J.K., K.H., K.K.; Supervision, K.K., K.H., O.W.W., S.J.K., C.G.; Funding Acquisition, K.K., K.H., O.W.W., S.J.K., C.G.

## Source data

Source Data File (Graf et al - Source Data.xlsx). The file contains all numerical data presented in Figures 1–6 and Tables S1–S6.

## Methods

### Key Resources Table

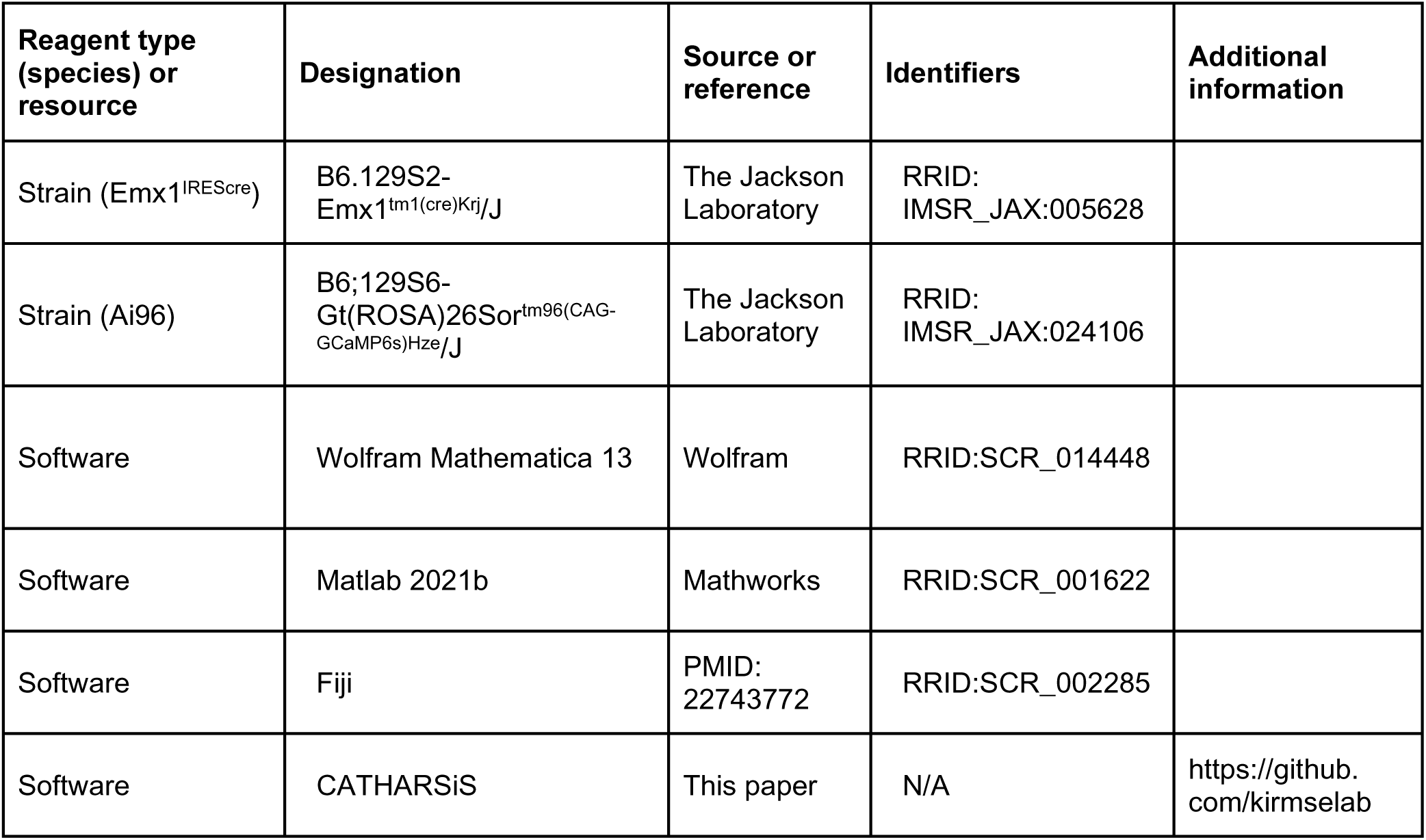

### Animals

All animal procedures were performed with approval of the local government (Thüringer Landesamt für Verbraucherschutz, Bad Langensalza, Germany; reference no.: 02-012/16) and complied with European Union norms (Directive 2010/63/EU). Animals were housed in standard cages with 14h/10h light/dark cycles. *Emx1^IREScre^* (stock no. 005628) and *GCaMP6s^LSL^* (Ai96, stock no. 024106) mice were originally obtained from The Jackson Laboratory. Double heterozygous offspring (*Emx1^IREScre/wt^:GCaMP6s^LSL/wt^*) was used for experiments at P3–4 (’P4’), P10–12 (’P11’) and P17–19 (’P18’). Mice of either sex were used.

### Surgical preparation, anesthesia and animal monitoring for in vivo imaging

30 minutes before starting the preparation, 200 mg/kg metamizol (Novacen) was subcutaneously injected for analgesia. Animals were then placed onto a warm platform and anesthetized with isoflurane (3.5% for induction, 1–2% for maintenance) in pure oxygen (flow rate: 1 l/min). The skin overlying the skull was disinfected and locally infiltrated with 2% lidocaine (s.c.) for local analgesia. Eyes of P17–19 were lubricated with a drop of eye ointment (Vitamycin). Scalp and periosteum were removed, and a custom-made plastic chamber with a central borehole (Ø 2.5–4 mm) was fixed on the skull using cyanoacrylate glue (Uhu) (P4: 3.5 mm rostral from lambda and 1.5 mm lateral from midline; P11: 3.5 mm rostral from lambda and 2 mm lateral from midline; P18: 3.5 mm rostral from lambda and 2.5 mm lateral from midline). For the hippocampal window preparation (Mizrahi et al., 2004), the plastic chamber was tightly connected to a preparation stage and subsequently perfused with warm artificial cerebrospinal fluid (ACSF) containing (in mM): 125 NaCl, 4 KCl, 25 NaHCO_3_, 1.25 NaH_2_PO_4_, 2 CaCl_2_, 1 MgCl_2_ and 10 glucose (pH 7.4, 35–36°C). A circular hole was drilled into the skull using a tissue punch (outer diameter 1.8 mm for P4 and 2.7 mm for P11 and P18 mice). The underlying cortical tissue and parts of corpus callosum were carefully removed by aspiration using a vacuum supply and a blunt 27G or 30G needle. Care was taken not to damage alveus fibers. As soon as bleeding stopped, the animal was transferred to the microscope stage.

During *in vivo* recordings, body temperature was continuously monitored and maintained at close to physiological values (36–37°C) by means of a heating pad and a temperature sensor placed below the animal. Spontaneous respiration was monitored using a differential pressure amplifier (Spirometer Pod and PowerLab 4/35, ADInstruments). Isoflurane was discontinued after completion of the surgical preparation and gradually substituted with the analgesic-sedative nitrous oxide (up to the fixed final N_2_O/O_2_ ratio of 3:1, flow rate: 1 l/min). Experiments started 60 min after withdrawal of isoflurane. At the end of each experiment, the animal was decapitated under deep isoflurane anesthesia.

In a separate set of experiments (Fig. 6), we analyzed the effects of N_2_O on animal state and CA1 network dynamics at P10–12. To this end, the following experimental timeline was applied in an additional cohort of six 6 animals (Fig. S3A). In the first FOV per mouse, Ca^2+^ imaging started under N_2_O (N_2_O/O_2_ ratio of 3:1 as above). About 10 minutes after replacing N_2_O by pure O_2_ (unanesthetized), Ca^2+^ imaging was continued in the same FOV (i.e. from the same cells). We then moved to a second FOV (with another set of cells) and performed recordings in a reversed order, i.e. Ca^2+^ imaging started under unanesthetized conditions before switching to N_2_O/O_2_ (recordings started 10 minutes after the onset of N_2_O administration). We reduced laser power and increased detector gain (as compared to recordings presented in Figs. 2–5) to prevent photo-bleaching in these longer-lasting experiments.

### Two photon Ca^2+^ imaging in vivo

After transferring the animal to the microscope stage, ACSF was removed and the hippocampal window was filled up with a droplet of agar (1%, in 0.9% NaCl) and covered with a cover glass. As soon as the agar solidified, the chamber was again perfused with ACSF.

Imaging was performed using a Movable Objective Microscope (Sutter Instrument) equipped with two galvanometric scan mirrors (6210H, MicroMax 673XX Dual Axis Servo Driver, Cambridge Technology) and a piezo focusing unit (P-725.4CD PIFOC, E-665.CR amplifier, Physik Instrumente) controlled by a custom-made software written in LabVIEW 2010 (National Instruments) (Kummer et al., 2015) and MPScope (Nguyen et al., 2006). Fluorescence excitation at 920 nm was provided by a tunable Ti:Sapphire laser (Chameleon Ultra II, Coherent) using a 20×/1.0 NA water immersion objective (XLUMPLFLN 20XW, Olympus). Emission light was separated from excitation light using a 670-nm dichroic mirror (670 DCXXR, Chroma Technology), short-pass filtered at 680 nm and detected by a photomultiplier tube (12 bit, H10770PA-40, Hamamatsu). Data were acquired using two synchronized data acquisition devices (NI 6110, NI 6711, National Instruments). Sampling rate was set to 11.63 Hz (256×256 pixels, 248×248 µm). For each animal, spontaneous activity was recorded from 3–5 FOVs, each one usually for ∼20 min. Some FOVs were excluded from further analysis due to excessive z-drifts. Finally, 1–4 FOVs were analyzed per animal and used for statistics. Care was taken to avoid any spatial overlap between sequentially recorded FOVs.

### Quantification and statistical analysis

#### Preprocessing

Image stacks were registered using NoRMCorre (Pnevmatikakis et al., 2017). For residual drift detection, a supporting metric was calculated as the Pearson correlation coefficient of the binarized template image used for stack registration and the binarized images of the registered image stack. Time periods with residual drift were then visually identified (by inspecting the supporting metric and the aligned image stack) and considered as missing values in subsequent analyses. Raw regions of interest (ROIs) were manually drawn around the somata of individual CA1 PCs using Fiji.

We quantified ΔF/F_0_ noise levels as the mean (per cell) difference between the 50^th^ and the 10^th^ percentile of the ΔF/F_0_ distribution. Noise levels were similar across the age groups (#5 in Table S2).

#### CATHARSiS – Calcium transient detection harnessing spatial similarity

For the detection of CaTs in densely labeled tissue, we devised CATHARSiS (<u>Ca</u>lcium transient detection <u>har</u>nessing <u>s</u>patial <u>s</u>imilarity). CATHARSiS makes use of the fact that spike-induced somatic GCaMP signals (ΔF) are spatially non-uniform and characteristic of a given cell. CATHARSiS comprises three major steps: (1) the generation of a spatial ΔF template representing the active cell, (2) the computation of a detection criterion *D(t)* for each time point (frame) and (3) the extraction of CaT onsets. All analyses were performed using custom scripts in Matlab and Fiji. CATHARSiS is available via GitHub (https://github.com/kirmselab).

Ad (1): For each ROI, we first obtained the mean F(t) by frame-wise averaging across all pixels of that ROI. We then computed the first derivative of F(t) and smoothed it using a second order Savitzky-Golay algorithm (window length, 6 frames), thus yielding Ḟ(t). We then determined eight candidate CaT onsets by extracting the frame numbers corresponding to the eight Ḟ(t) peaks having the largest amplitude. This step was performed in an iterative-descending manner by starting with the largest F(t) peak. For each peak, we defined a minimum time difference (5 frames) to all subsequently extracted peaks, so as to avoid extracting nearby frames belonging to the same CaT. For each candidate CaT onset, we then computed the corresponding spatial ΔF (average of five successive frames). To this end, we first radially expanded the raw ROI by two pixels using the Euclidian distance transform (we found that this increased detection reliability due to enhanced spatial contrast). Resting fluorescence F_0_(t) was defined as the moving median over 500 frames. Eight candidate ΔF templates were obtained by converting raw ΔF values into z-scores. Based on visual inspection, we next rejected those candidate ΔF templates that putatively reflected activation of optically overlapping somata and/or neurites. If all candidate ΔF templates had been rejected, the cell was excluded from further analysis; otherwise, the remaining candidate ΔF templates were averaged to obtain the final ΔF template representing the active cell.

Ad (2): For each ROI (spatially expanded as above), we extracted its spatial ΔF for all frames in the image stack. Next, the spatial ΔF template representing the active cell was optimally scaled to fit its ΔF in each recorded frame. Based on the optimum scaling factor and the goodness of the fit, a detection criterion *D(t)* was computed for each time point. Here, *D(t)* was defined without modification as previously described for the temporal domain (Clements et al., 1997).

Ad (3): For each ROI, CaT onsets were extracted from *D(t)* using UFARSA, a general-purpose event detection routine (Rahmati et al., 2018). To this end, we slightly modified the original UFARSA approach in two ways. I) Following the smoothing step implemented in UFARSA, all negative values were set to zero, as we found in our perliminary analysis that negative-to-positive transitions occasionally resulted in false positive events. II) We introduced a lower bound for the leading threshold, so as to minimze potential false positive events. Reconstructed CaT onsets were translated into a binary activity vector (1 – event, 0 – no event) and used for the following analyses.

#### Firing irregularity

For each cell, we quantified the irregularity of its CaT onsets (i.e. firing times) using CV2, as a local and relatively rate-independent measure of spike time irregularity (Holt et al., 1996; Ponce-Alvarez et al., 2010): 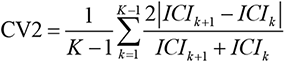, where *ICI_k_* and *ICI_k+1_* are the *k*th and *(k+1)*th inter-CaT intervals of the cell, and *K* is the total number of its *ICI* s. To achieve more robust results cells with less than ten *ICI* s were excluded from this analysis.

#### Network bursts

Network bursts (NBs) were defined as a significant co-activation of cells as follows: (1) To account for some temporal jitter in the detection of CaT onsets, all values in the binary activity vectors that fell within ±Δt frames of any detected CaT were set to 1. Unless otherwise stated, Δt was set to 3. We then computed the mean across the resulting activity vectors of all individual cells to obtain the empirical fraction of active cells per frame Φ(t). (2) We randomly shuffled CaT onsets of all cells (uniform distribution; 1,000 times), computed the surrogate Φ(t) (as above) and defined the 99.99th percentile of all surrogate Φ(t) as the threshold for NB detection. The NB threshold was determined separately for each FOV, so as to account for different mean CaT frequencies. (3) Any frame with an empirical Φ(t) exceeding the threshold was considered as belonging to an NB.

To examine the robustness of our findings, we systematically varied the operational definition of the threshold used for NB detection in two ways. (1) In the first approach, Δt was set to values ranging from 1 to 11 frames. A frequency-dependent threshold was then computed for each FOV as detailed above (Fig. S1A). (2) In the second approach, a constant (frequency-independent) threshold was applied to all FOVs (ranging from 7 to 17% active cells per frame) (Fig. S1B). Here, Δt was set to 3 frames. Note that threshold values below ∼10% are less meaningful, as the average fraction of active cells in some FOVs at P18 is ∼9%.

In the resulting binary NB vectors, 0-1 transitions were defined as NB onsets and 1-0 transitions as NB offsets. Using the binary NB vectors, we extracted (i) the relative time the network spent in NBs and (ii) the average NB duration. NB size was defined as the fraction of cells which were active in at least one frame of a given NB, corrected for the chance level of co-activation by subtracting the NB threshold.

#### Discontinuous and continuous network activity

We partitioned recordings into non-overlapping time bins of 116 frames (∼10 s) duration. Network activity during a given time bin was classified as continuous if the fraction of active cells per frame Φ(t) exceeded 3% in >70% of all frames belonging to that bin; if Φ(t) exceeded 3% in ≤70% of all frames belonging to that bin, it was considered discontinuous. To compute Φ(t), Δt was set to 3 frames.

#### Power analysis

To account for missing values representing the residual drift periods (see above), spectral power of the fraction of active cells Φ(t) was estimated by computing the Lomb-Scargle periodogram (Matlab, MathWorks). To compute Φ(t), Δt was set to 3 frames.

#### Pairwise correlations

Spike-time tiling coefficients (STTCs) were computed for all possible cell pairs with a synchronicity window Δt of three frames (∼258 ms) using custom written code (Matlab, MathWorks) (Cutts et al., 2014). STTCs derived from measured data were compared to those from surrogate data obtained by randomly shuffling (uniform distribution; 1,000 times) CaT onsets of all cells, separately. This randomization kept the mean CaT frequency of each cell unchanged.

#### Population coupling

To quantify the degree of coupling of each cell to the overall population activity we computed its population coupling, PopC (Okun et al., 2015; Sweeney et al., 2020). To this end, for each cell, we first smoothed its binary vector (see above) and the summed vector of the rest of population, followed by computing PopC as the Pearson correlation coefficient between these two vectors. For smoothing, we used a Gaussian kernel with *SD* = 3 frames. To assess the significance of the PopCs (i.e. being beyond chance) we generated surrogate data by binning the raster matrix along time-axis; non-overlapping bins with a size of 10 frames (ca. *SD*, according to (Kruskal et al., 2007)). We randomly exchanged CaT onsets across active cells within each bin (500 times), thereby effectively preserving the CaT frequency of each cell as well as the local summed activity of the population. For each cell, using its surrogates, we determined the significance of its empirical PopC (95^th^ percentile). Moreover, when reporting the PopC of each cell, we subtracted the mean of its surrogate PopCs, in order to account for the potential differences in population activity levels of different FOVs (for a similar approach see (Okun et al., 2015; Sweeney et al., 2020)). Cells with less than five CaTs were excluded from this analysis, to increase robustness of our results.

#### Motifs of population activity

To identify the specific cellular activation patterns recurring over time (i.e. motifs of population activity) we used an eigendecomposition-based clustering method (Li et al., 2010; Patel et al., 2015). To this end, we first divided the recording time into non-overlapping windows with a size of 10 frames, and assigned 1 and 0 to cells which were active or silent during each bin. This converts the raster matrix to a sequence of binary vectors (i.e. spatial patterns), where each pattern has a size of Nx1 (N is the number of analyzed cells in the FOV). We then computed the degree of similarity between all possible pairs of these patterns using matching index (Romano et al., 2015): 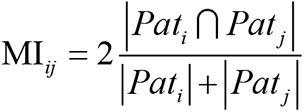, *Pat_i_* and *Pat_j_* are the *i*th and *j*th binary cellular activation patterns (vectors), and the norms are equal to the number of ones (i.e. active cells) in each vector. MI ranges from 0 (no similarity) to 1 (perfect similarity), and in particular approximates the number of common neuronal activations (i.e. common ones) between pattern pairs; for more details see (Romano et al., 2015; Sporns et al., 2007). Accordingly, for each FOV, we obtained a similarity matrix of size P x P, where P indicates the number of patterns. The rows and columns relating to the silent-pattern pairs were excluded, as they were giving rise to an undefined value (i.e. 0 divided by 0). We used the MI matrix as the input to the eigendecomposition clustering method. Briefly, this method decomposes a given similarity matrix (here, MI matrix) into a set of eigenvalues and eigenvectors: The number of significantly large eigenvalues determines the number of motifs, and their corresponding eigenvectors contain the information about motif structure (i.e. the set of patterns belonging to each motif). The largest eigenvalue is proportional to the global similarity among all patterns. As the surrogate data for testing the statistical significance of the eigenvalues and also computing a normalized unbiased value of global similarity index, we used the randomly shuffled CaT onsets (see above), based on which we repeated the binning and computation of MI matrices (500 times). This procedure enabled us to identify the motifs of cellular activation patterns, which occurred beyond chance level. For more details about the clustering method and its mathematical description see (Li et al., 2010).

#### Analysis of cardiovascular parameters and movement periods

Respiration and movement were recorded by means of an air pillow positioned below the chest of the animal and a differential pressure amplifier (Spirometer Pod and PowerLab 4/35, ADInstruments). We first computed the short-time Fourier transform using a time window of one second with an overlap of 50% (Fig. S2). To extract respiration and heart rates, the median power spectral density was calculated across time, and the first two peaks were detected. In one FOV this was not possible, as the two peaks could not be reliably separated. For the detection of movement periods, bandpower (0.1–8 Hz) was first calculated across time. A time bin was classified as movement, if the bandpower exceeded a threshold, defined as the moving median (over 60 s) plus three times the moving absolute deviation (over 300 s). In the resulting binary movement vectors, 0-1 transitions were defined as movement onsets and 1-0 transitions as movement offsets.

### Computational modeling of a developing neural network with inhibitory GABA

#### Overview

To gain insights into the mechanisms and functional role of the observed network burstiness during the emergence of synaptic inhibition in CA1, we used computational modeling and stability analysis. For this purpose, we employed a recently established model of a recurrent neural network (RNN) for first postnatal month development (Rahmati et al., 2017). It is an extended Wilson-Cowan-type model (Tsodyks et al., 1998), and benefits from being biophysically interpretable and mathematically accessible. Recently, this model was also adapted successfully to explain key dynamics and mechanisms of GDPs in neonatal CA1 with excitatory GABA signaling during the first postnatal week (Flossmann et al., 2019). However, in accordance with previous reports and our present experimental data for the second postnatal week, we here use the model with mainly two specific cellular properties: I) GABAergic synapses are considered inhibitory (Kirmse et al., 2015; Murata et al., 2020; Valeeva et al., 2016), and II) the mean spontaneous firing activity of PCs is effectively non-zero (Fig. 2C). In the following, after providing the mathematical description of the model, we describe the mathematical components used for its stability analysis. For more details about the model and the approach see (Rahmati et al., 2017).

#### Model description

The model is a mean-field network model of mean firing activity rates of two spatially localized, homogeneous glutamatergic and GABAergic cells (here, pyramidal (PC) and interneuron (IN) populations) that are recurrently connected (Fig. 7A). The model incorporates two short-term synaptic plasticity (STP) mechanisms, namely short-term synaptic depression (STD) and facilitation (STF), which render the synaptic efficacies dynamic over time. Hence, we call the network hereafter STP-RNN. The equations governing the mean-field dynamics of the STP-RNN (10D) are (dots denote the time derivatives and, hereafter, PC and IN are abbreviated as P and I for readability) (Rahmati et al., 2017):

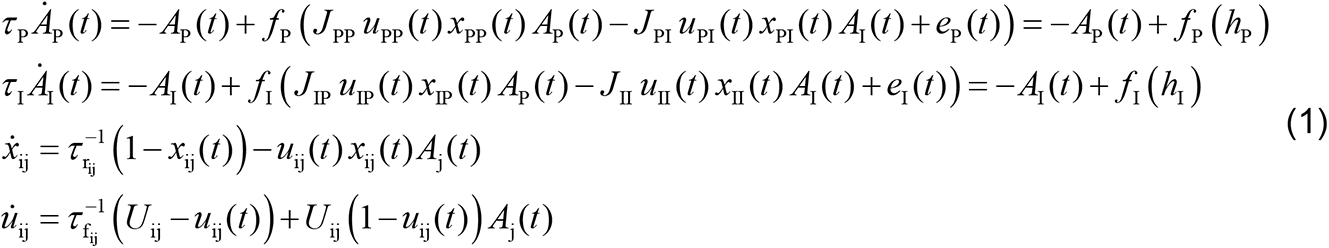

where i and j∈{P,I}, and j is the index of the presynaptic population, *A*_P_ and *A*_I_ are the average activity rates (in Hz) of PC and IN populations which can be properly scaled to represent locally the average recorded activities in these populations, *x*_ij_ and *u*_ij_ are the average dynamic variables of STD and STF mechanisms, *τ*_P_ and *τ* _I_ are approximations to the decay time constants of the glutamatergic and GABAergic postsynaptic potentials, *τ*_ij_ is the synaptic recovery time constant of depression, *τ*_ij_ is the synaptic facilitation time constant, *U*_ij_ is analogous to the synaptic release probability, *J*_ij_ is the average maximum absolute synaptic efficacy of recurrent (i=j) or feedback (i ≠ j) connections, and *e*_P_ and *e*_I_ are the external inputs received by the PC and IN populations from other brain regions or stimulation. In this work, we set the inputs to zero (for spontaneous baseline activity), or model them as excitatory pulse (with variable positive amplitude) with a duration of 20 ms thereby emulating e.g. the SPW-driven inputs to the PC and IN populations (Karlsson et al., 2006). The transformation from the summed input to each population, *h*_i_, to an activity output (in Hz) is governed by the response function, *f*_i_, defined as:

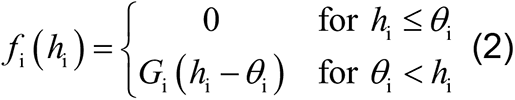

where *θ*_i_ is the population activity threshold, and *G*_i_ is the linear input-output gain above *θ*_i_. In this work, we parameterize the STP-RNN as a network model representing mainly a stage during the second postnatal week. To do this, we mainly followed (Rahmati et al., 2017) by setting 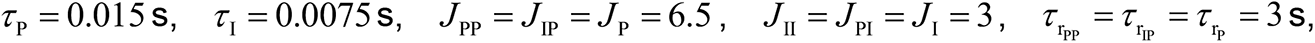 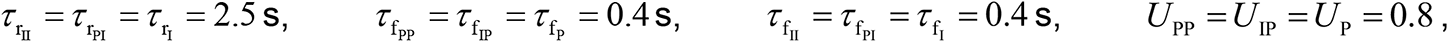 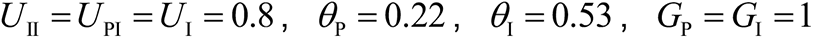 and *e*_P_ = *e*_I_ = 0 Hz (for spontaneous baseline activity). According to these parameter values: I) both glutamatergic and GABAergic connections will act depressing, and II) the network will spontaneously have, in addition to a silent state, an active state where both *A*_I_ and, in particular, *A*_P_ are effectively non-zero, and III) GABAergic transmission will be inhibitory (note the positive value of *J*_I_). Note that points II) and III) render the model inherently different from the neonatal STP-RNN used by (Flossmann et al., 2019).

#### Frozen STP-RNN

A Frozen STP-RNN is obtained by freezing the synaptic efficacies of a STP-RNN; i.e. by fixing the STP variables *x*_ij_ and *u*_ij_ at the values of interest. This will convert the STP-RNN (10D; see Eq. 1) effectively to a 2D network with constant synaptic weights. As shown in (Rahmati et al., 2017) and (Flossmann et al., 2019), the Frozen STP-RNN can provide a reliable approximation to the stability behavior of a STP-RNN at the state chosen for freezing (see below). The equations governing the dynamics of a Frozen STP-RNN are:

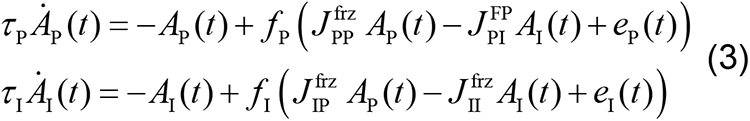

where 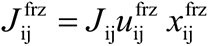 and 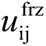 and 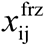 are the values of *u*_ij_ and *x*_ij_ (see Eq. 1) at the state of interest; here, at a silent state, active state, or the time of network burst’s peak (see Results). *Phase plane.* To visualize the stability behavior of our network model, we used the phase plane analysis based on the activity rates: *A*_I_ -*A*_P_ –plane (2D). The *A*_I_ -*A*_P_ –plane sketch includes the curves of the *A*_P_ –nullcline and *A*_I_ –nullcline representing sets of points for which *A*_P_ (*t*) = 0 and *A*_I_ (*t*) = 0. Any intersection of these nullclines is called a fixed point (FP), with the stability needed to be determined (see below). For the STP-RNN, these FPs represent the steady states of the full network, i.e. the 10D STP-RNN in Eq. 1 (see also Fig. 7B). For the Frozen STP-RNN (thus, 2D; see Eq. 3) with synaptic efficacies frozen at the state of interest (e.g. silent state), these FPs may include that state, and possibly some other FPs which may not exist in the STP-RNN itself (e.g. see Figs. 7D and 8C). In addition to the visualization of the FPs in the *A*_I_ -*A*_P_-plane, we also computed the FPs by numerically solving Eq. 1 and Eq. 3 (separately) after setting the right hand side of the equations to zero. For more details see (Rahmati et al., 2017).

#### Stability of FPs

To determine the stability of any FP in the STP-RNN (resp. in the Frozen STP-RNN) we applied the linear stability analysis to its 10D (resp. 2D) system of equations in Eq. 1 (resp. Eq. 3): We investigated whether all eigenvalues of the corresponding Jacobian matrix have strictly negative real parts (if so, the FP is stable), or whether at least one eigenvalue with a positive real part exists (if so, the FP is unstable).

#### Simulations

All simulation results in this paper have been implemented as Mathematica and Matlab (MathWorks) code. For network simulations, we set the integration time-step size to 0.0002 s. In Figs. 7C and S5B–E, the initial conditions of the STP-RNN variables were set to those values of the spontaneous stable FP of the network at the active state.

#### Operating regimes and FP-domains

The stable operating regimes of a RNN at a FP can be classified as an inhibition-stabilized network (ISN) vs. a Non-ISN (Latham et al., 2004; Ozeki et al., 2009; Rahmati et al., 2017; Tsodyks et al., 1997). To apply this theoretical classification to the STP-RNN model, we used the previously described analytical findings and numerical techniques (for details see Rahmati et al., 2017). In brief, to discriminate between these two regimes in STP-RNN with inhibitory GABAergic synapses three criteria were defined: (A) Excitatory instability: For the inhibitory activity rate fixed at the FP, the recurrent excitation is strong enough to render the PC-population intrinsically unstable. (B) Excitatory stability: In contrast to (A), the PC-population is stable *per se*, i.e. even with a feedback inhibition fixed at its level at the FP. (C) Overall stability: The dynamic feedback inhibition to the PC-population is strong enough to stabilize the whole network activity. At a FP, a network operating under the (A) and (C) criteria is an ISN, while a network operating under the (B) and (C) criteria is a Non-ISN. A network, which is neither ISN nor Non-ISN at the FP, operates under an unstable regime. Clearly, for the network with excitatory GABAergic synapses, the ISN regime cannot be defined. Therefore, at a FP, the network is either unstable or Non-ISN. However, as in this case the condition (C) is not applicable, the Non-ISN refers to a non-unstable regime; i.e. where all eigenvalues of the corresponding Jacobian matrix at the FP have strictly negative real parts (thus, FP is stable).

In this framework, the *A*_I_ -*A*_P_-plane is partitioned into different domains of operating regimes (FP-domains). Each FP-domain contains all potential steady states (i.e. FPs) at which the network could operate under the corresponding regime. The FP-domains of operating regimes were determined by using numerical simulations, based on the aforementioned stability criteria obtained analytically. The area of each regime’s FP-domain is computed numerically using a sparse grid rule as implemented in Mathematica (version 13).

#### Alternative network models

To assess whether, or to what extent, our observed CA1 dynamics in second postnatal week can be explained by other network models (or mechanisms), we created two operationally distinct network models by reparametrizing the STP-RNN model (Eq. 1). (I) Mono-RNNi (θ_P_ ↓, ‘i’ for inhibitory GABA): θ_P_ = −0.18. (II) Mono-RNNe: (J_I_ → −0.5×J_I_, θ_P_ ↓, θ_I_ ↓; ‘e’ for excitatory GABA): θ_P_ = −0.3, θ_I_ = −0.1, J_I_ = −1.5. Either of models (10D) has only one spontaneous FP which is stable (thus, mono-stable) and located at an active state. The properties of these models have been detailed in Results (corresponding text of Fig. S5). According to these parameter values, GABAergic transmission in Mono-RNNe is excitatory rather than inhibitory (note the negative value of its J_I_). However, it has a weaker excitatory effect than glutamatergic transmission; i.e. |J_I_|< (|J_P_|= 6.5). Moreover, note that the lower population activity-threshold (θ) of each population reflects that its neurons receive a higher mean level of spontaneous background input and/or have a higher intrinsic excitability (e.g. lower spike-threshold/rheobase or higher membrane resistance, see also Flossmann et al., 2019; Rahmati et al., 2017).

### Statistical analysis

Statistical analyses were performed using OriginPro 2018 and Microsoft Excel 2010 using the Real Statistics Resource Pack software (Release 7.2, Charles Zaiontz). Unless otherwise stated, the statistical parameter n refers to the number of FOVs (P4: 19 FOVs from six animals, P11: 11 FOVs from six animals, P18: 12 FOVs from six animals). All data are reported as mean ± standard error of the mean (SEM), if not stated otherwise. The Shapiro–Wilk test was used to test for normality. Homogeneity of variances was tested with the Levene’s test using the median. For multi-group comparisons, analysis of variance (ANOVA) was applied for normally distributed data or the Kruskal-Wallis test for non-normally distributed data. In the case of unequal group variances, Welch’s correction was applied for the ANOVA. Following a significant result in the ANOVA, *post-hoc* pairwise comparisons were performed using the Tukey-Kramer (equal variances) or the Games-Howell (unequal variances) test. Following a significant result in the Kruskal-Wallis test, *post-hoc* pairwise Mann-Whitney U-tests following Holm’s approach were performed. P values (two-tailed tests) < 0.05 were considered statistically significant, except for the Shapiro-Wilk test (P < 0.01). Details of the statistical tests applied are provided in Tables S1–S6.

### Data and code availability

All data analyzed during this study are included in the manuscript, Supplementary Information and supporting Source Data File. CATHARSiS is available via GitHub (https://github.com/kirmselab).

