## Supplementary Information for "Network instability dynamics drive a transient bursting period in the developing hippocampus *in vivo*"

**This file includes:**

Supplementary Text
Figures S1 to S5
Tables S1 to S6

### Supplementary Text

*Facing a new deadline once the transition to the active state has failed*

Here, we address how the network can transition to the active state once a simNB failed to converge to the active state (thus, the network returned to the silent state). This can occur in two cases: 1) if the input is relatively strong (see the light green area on the right side of Fig. 7J), and 2) if the internal deadline is missed (Fig. 8G). Here, we address this question for the latter case, while our findings will be similarly applicable to the former one.

To this end, we first introduce a third input pulse arriving after the second one whose evoked simNB failed to push the network to the active state (Fig. S4A and D); recall that the first input pulse was used for silencing the network (see Fig. 7C). We found that in this case the network encounters a new deadline (Fig. S4A and S4D; dotted black line #2). In addition, the network expresses a refractory period after the first simNB: Any input (regardless of its strength) arriving during the refractory period will not be able to move the network to the active state, and may not be able to trigger a simNB. This results from a weakening of synaptic weights by the first simNB (Rahmati et al., 2017), precluding the network from forming the required transient unstable (allowing for simNB emergence) and stable (allowing for transitioning to the active state) FPs in its fast dynamics. Once the network recovers sufficiently to generate a simNB (Fig. S4 B, C, E, F), the countdown for the arrival of a third input to initiate the transition begins (Fig. S4G). As compared to the second input, the third input – with a proper ratio – has a shorter time-window to enable the transition (compare Figs. 8G and S4G). This is mainly because of the emerged refractory period. Note that, in contrast to the deadlines, the refractory period is not fixed but has a direct relationship to the size of the preceding simNB; a larger simNB (returning to the rest state) will result in a longer refractory period, which is needed for sufficient synaptic recovery (Rahmati et al., 2017). Except for the refractory period, the rest of the mechanisms and responses of the network remain similar to the case of the first deadline (see the corresponding text of Fig. 8).

Collectively, these results indicate that, in addition to the input ratio, a delicate interaction between the input timing and the network internal dynamics associates with CA1 input-encoding schemes prior to the onset of environmental exploration.

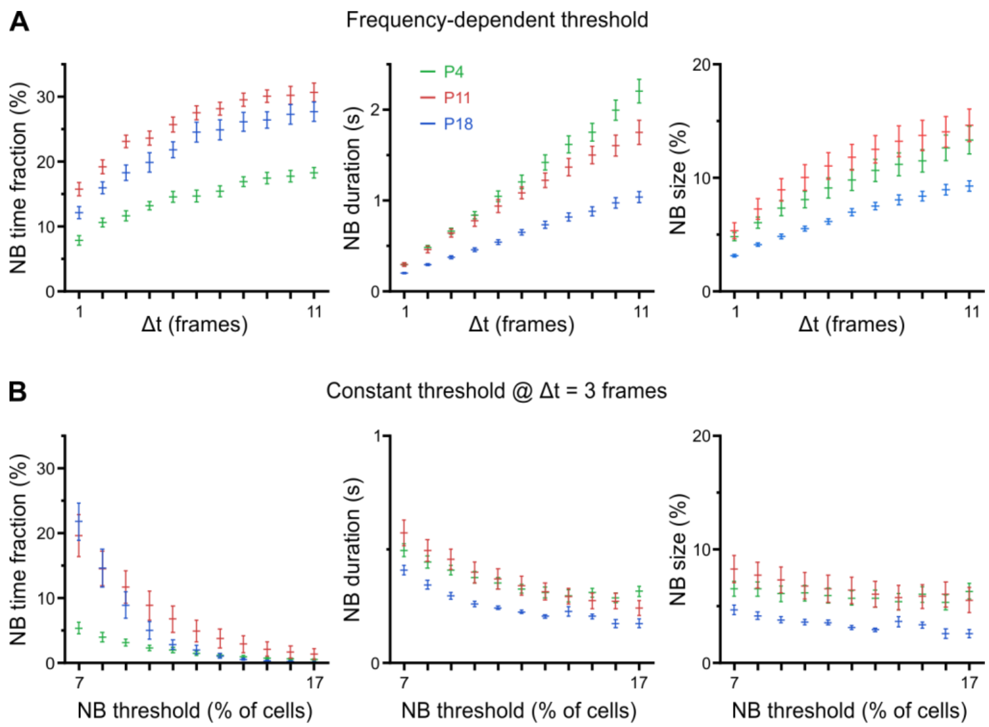

**Figure S1. Related to Figure 3.** Developmental changes in network burst (NB) characteristics are robust to a wide range of definitions. (A) Total time spent in NBs (*left*), mean NB duration (*middle*) and mean NB size (*right*) as a function of the parameter Δt. Δt is used for computing a CaT frequency-dependent threshold, separately for each FOV (see Methods for details). Sampling rate was 11.63 Hz. (B) Total time spent in NBs (*left*), mean NB duration (*middle*) and mean NB size (*right*) as a function of a constant threshold applied to all FOVs, i.e. independent of CaT frequencies. Note that threshold values below ~10% are less meaningful, as the average fraction of active cells in some FOVs at P18 is ~9%. Δt was set to 3.

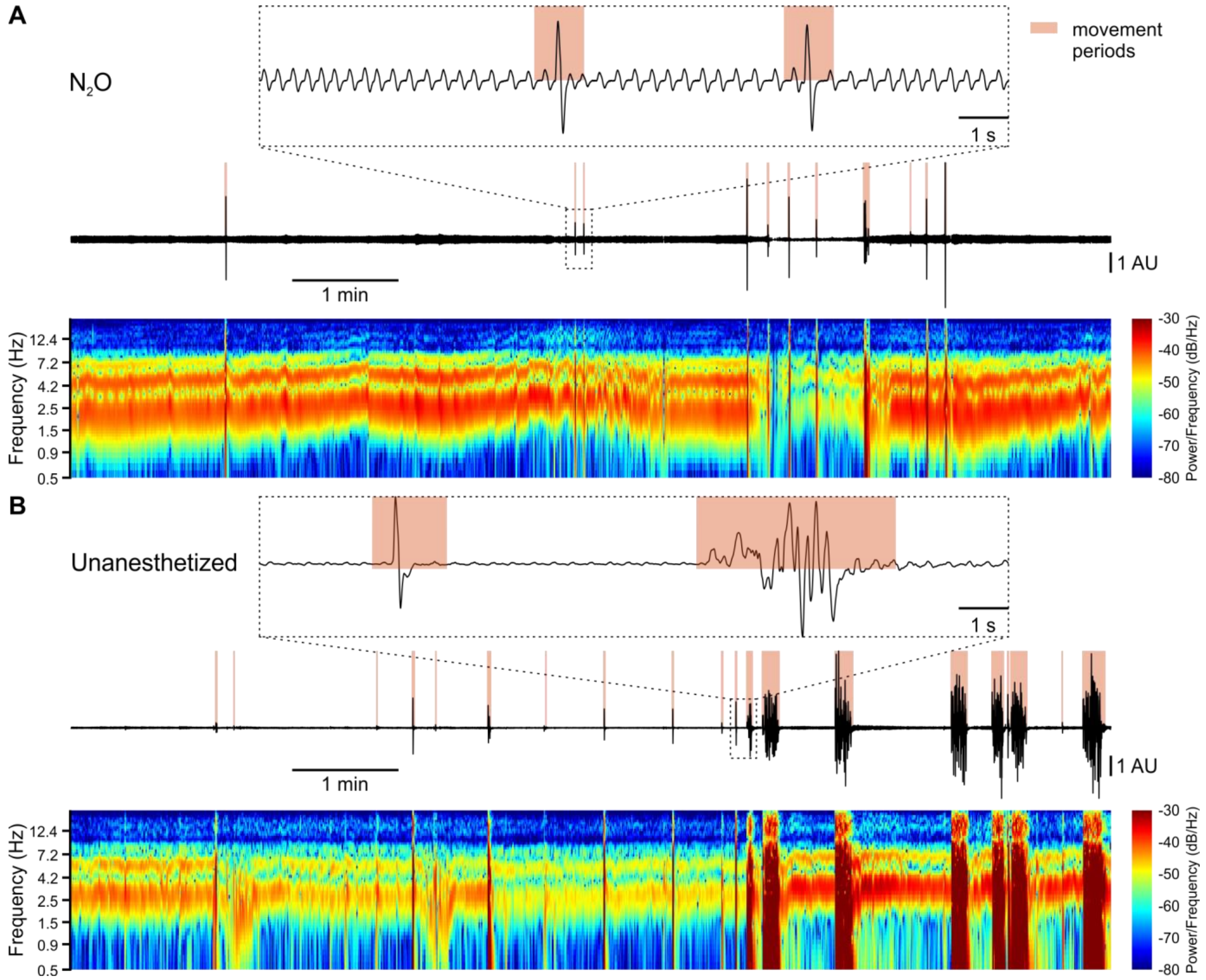

**Figure S2. Related to Figure 6.** Detection of body movements, breathing and heart rate. (A and B) Sample respiration/movement signal (*middle*) and time-aligned spectrogram (*bottom*; 0.5–20 Hz, window length: 1s, overlap: 50%) used to detect movement periods, respiration and heart rate. *Top*: Marked time periods (dotted rectangle) at higher temporal resolution. Signals were recorded by means of a pressure sensor positioned below the chest of the animal. (A) Recording from an animal receiving 75% $N_2O$/25% $O_2$ ('$N_2O$'). (B) Recording from the same mouse receiving pure oxygen ('Unanesthetized').

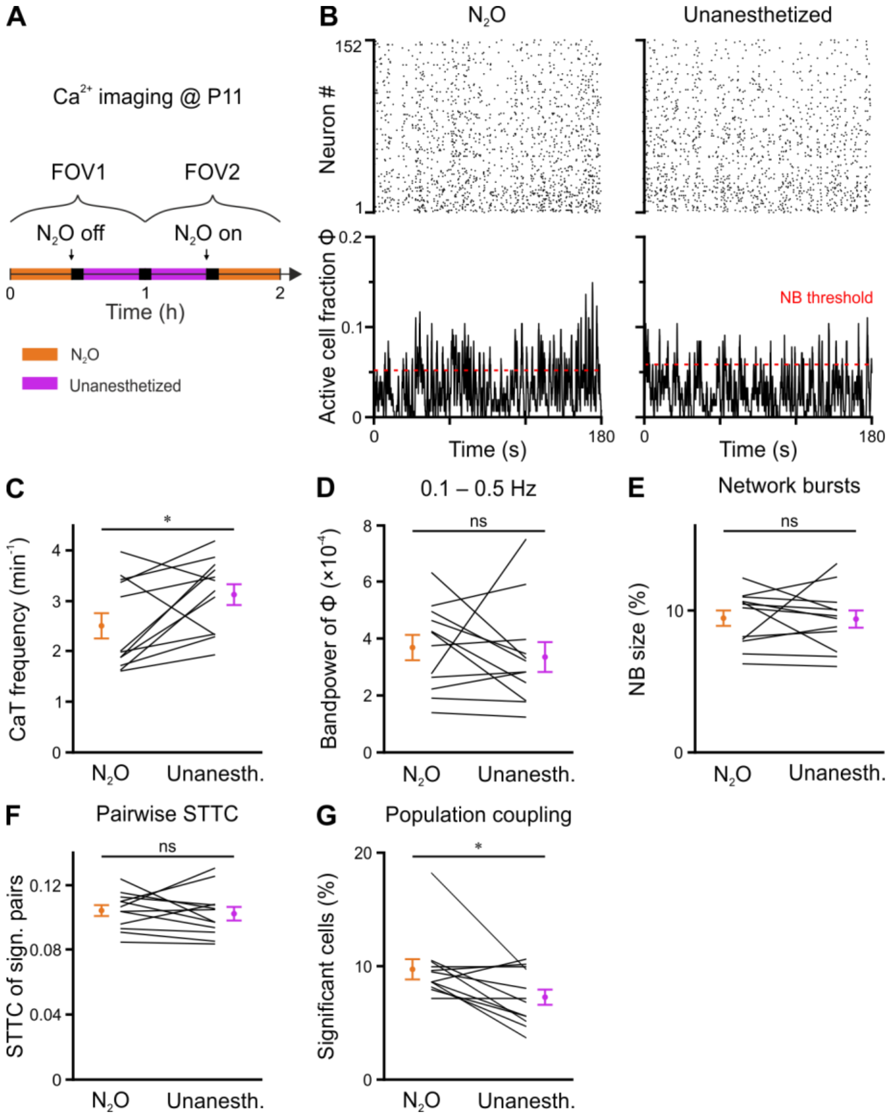

**Figure S3. Related to Figure 6.** Effects of nitrous oxide on CA1 network dynamics at P11. (A) Experimental timeline. In each animal, two FOVs were recorded with and without $N_2O$ (paired design). (B) Rasterplots (*top*) and time-aligned fraction of active cells Φ(t) (*bottom*) from the same FOV in either the presence or the absence of $N_2O$. Red dotted lines indicate the activity-dependent thresholds for NB detection. (C) Mean CaT frequency per FOV. (D) Bandpower of Φ(t) in the 0.1–0.5 Hz range. (E) NB size quantified as the mean fraction of active neurons per NB (corrected for burst threshold as indicated in B). (F) Mean STTC of significantly correlated cell pairs. (G) Mean fraction of cells with significant population coupling (PopC). n = 12 FOVs from six mice, * $P < 0.05$, ns – not significant. See also Table S6 and Source Data File.

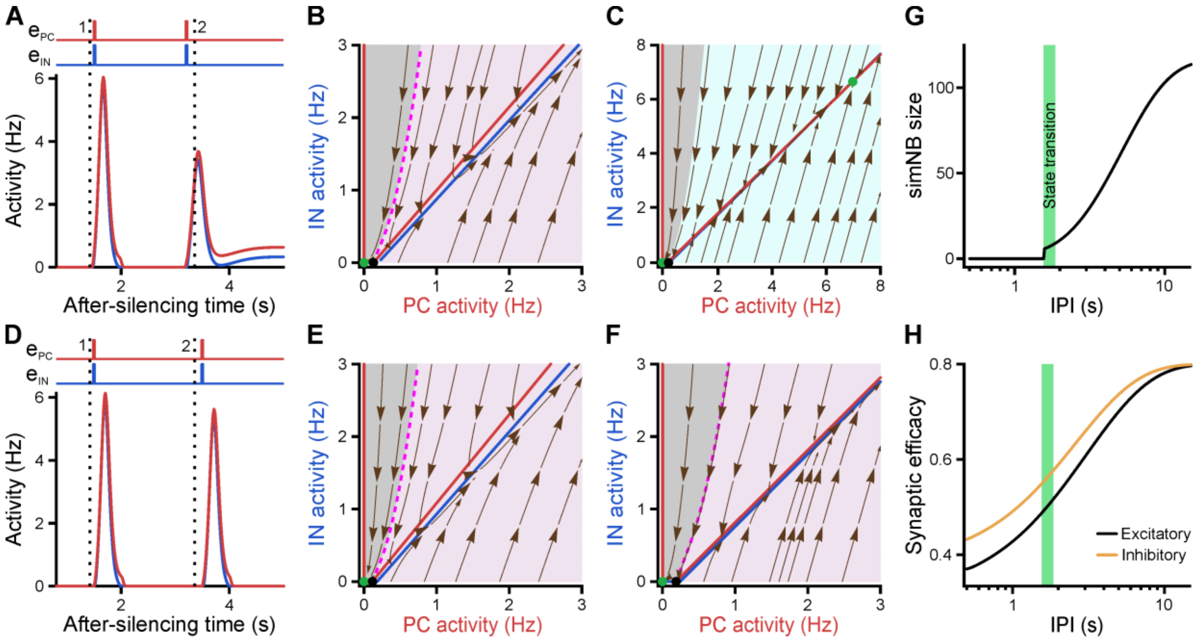

**Figure S4. Related to Figure 8.** Re-emergence of an internal deadline after a transition failure. (A-H) Same format as in Fig. 8. Following the failure of the network in transitioning to the active state due to missing the deadline (dotted black line #1, at t = 1.45 s), the subsequent input delivered to the network faces a new deadline (#2, at t = 3.36 s). $e_{\mathrm{PC}} = 0.25$, $e_{\mathrm{IN}} = 0.25$. (A–C) Input delivered to the network before the new deadline (#2) can move it to the active state. Inputs delivered at t = 1.55 s and t = 3.2 s, relative to the time that the network was silenced (see Fig. 7F). (D–F) Once the new deadline (#2) is missed, the network fails to transition to the active state by the input. Inputs delivered at t = 1.55 s and t = 3.5 s. (G) The size of the simNB and the network transition to the active state depend on the IPI. (H) Same as G, but for the non-scaled efficacies of GABAergic and glutamatergic synapses, right before the arrival of the third input. The first input pulse used for silencing the network is not shown.

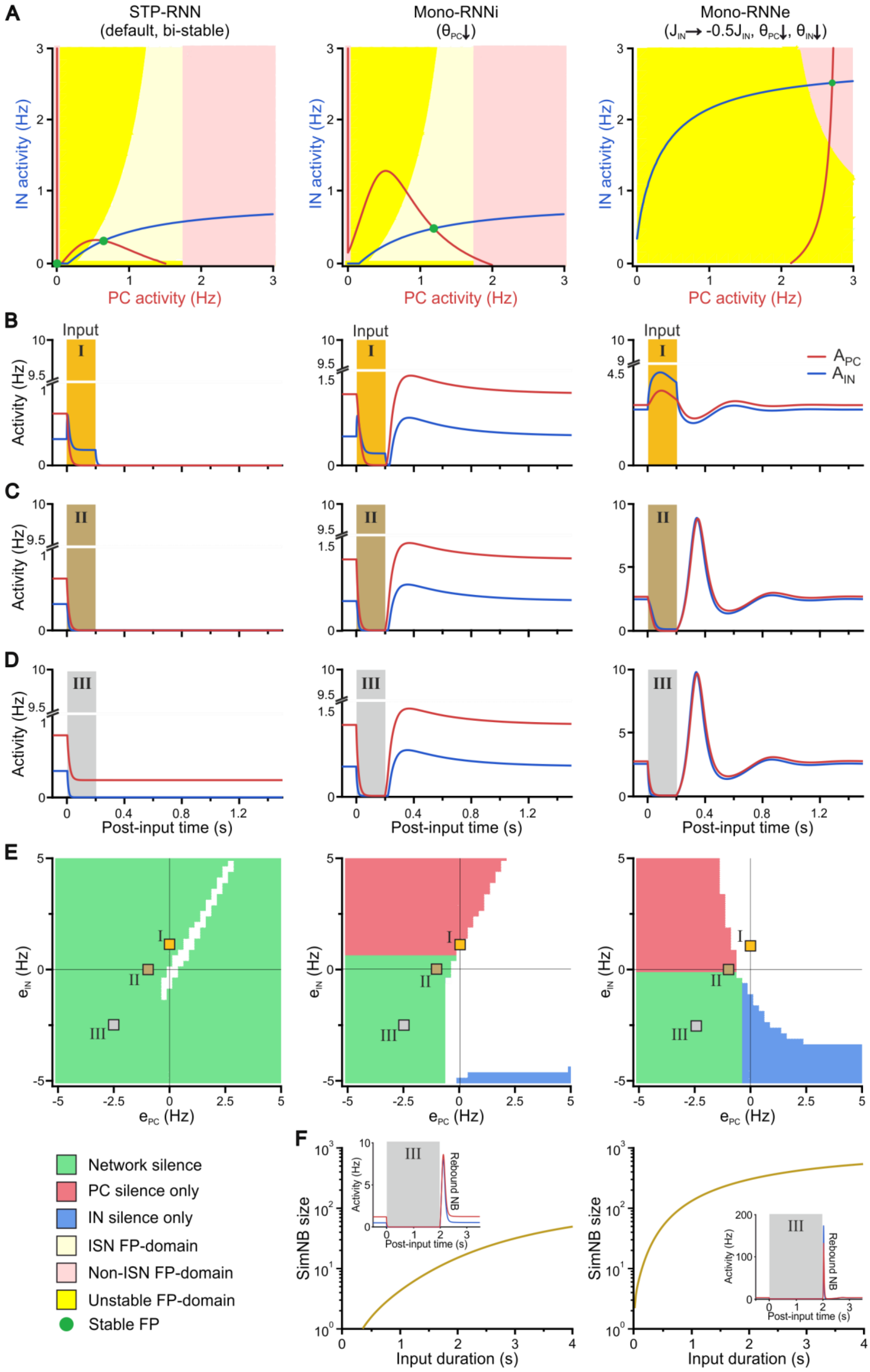

A
STP-RNN
(default, bi-stable)
Mono-RNNi
(θPC↓)
Mono-RNNe
(JIN→ -0.5JIN, θPC↓, θIN↓)
IN activity (Hz)
PC activity (Hz)
B
Input
I
Activity (Hz)
APC
AIN
C
II
D
III
Post-input time (s)
E
eIN (Hz)
ePC (Hz)
Network silence
PC silence only
IN silence only
ISN FP-domain
Non-ISN FP-domain
Unstable FP-domain
Stable FP
F
SimNB size
Input duration (s)
Rebound NB

**Figure S5. Related to Figures 8 and 9.** A bi-stable STP-RNN model with inhibitory GABA robustly explains the experimental observations. (A) In contrast to the bi-stable STP-RNN, Mono-RNNi and Mono-RNNe networks lack a silent state (i.e. a FP at origin). Same format as Fig. 9A, overlaid by the $A_{\mathrm{PC}}$-nullcline (red), the $A_{\mathrm{IN}}$-nullcline (blue) and the FPs of each network with dynamic synapses (i.e. 10D system). The unstable FP in the STP-RNN is not shown for clarity. For the exact values of model parameters, see Methods. (B–C) Network responses to example inputs (200 ms; shaded areas). In contrast to the STP-RNN, Mono-RNNi and Mono-RNNe networks can be silenced only during the input. Inputs are: $e_{\mathrm{PC}}=0$, $e_{\mathrm{IN}}=1$ (B); $e_{\mathrm{PC}}=-1$, $e_{\mathrm{IN}}=0$ (C); $e_{\mathrm{PC}}=e_{\mathrm{IN}}=-2.5$ (D). (E) In contrast to the STP-RNN, silencing of Mono-RNNi and Mono-RNNe models requires the input (200 ms) to at least one of network populations to be inhibitory. Green indicates complete network silence ($A_{\mathrm{PC}}=A_{\mathrm{IN}}=0$), red indicates PC silence only ($A_{\mathrm{PC}}=0$, $A_{\mathrm{IN}}\neq 0$), and blue indicates IN silence only ($A_{\mathrm{PC}}\neq 0$, $A_{\mathrm{IN}}=0$). White areas: $A_{\mathrm{PC}}\neq 0$, $A_{\mathrm{IN}}\neq 0$. The activity levels of the neuronal populations were computed as those prior to the input removal. Square symbols (I–III) indicate the input combinations used in panels B–D. (F) In contrast to the STP-RNN, effective (rebound) simNB generation in Mono-RNNi (*middle*) and Mono-RNNe (*right*) requires a relatively long-lasting input. (Inset) Same as D, but for a longer input of 2 seconds.

**Table S1.** Synopsis of statistical tests related to Figure 1. Numerical data are provided in the Source Data File.

| # | Related to | Trace | Descriptive statistics | N | Test | Test statistics |
|---|---|---|---|---|---|---|
| | Figure 1I | | | | | |
| 1 | Recall | D | 0.953 ± 0.017 | 20 | Wilcoxon Signed-Rank Test | P = 0.44<br>W = 57<br>Z = 0.77 |
| | | ΔF | 0.947 ± 0.014 | 20 | | |
| 2 | Precision | D | 0.915 ± 0.015 | 20 | paired t-test | $P = 1.4 \times 10^{-6}$<br>t = 6.9<br>df = 19 |
| | | ΔF | 0.607 ± 0.047 | 20 | | |
| 3 | F1 | D | 0.930 ± 0.010 | 20 | paired t-test | $P = 8.0 \times 10^{-6}$<br>t = 6.1<br>df = 19 |
| | | ΔF | 0.719 ± 0.037 | 20 | | |

**Table S2.** Synopsis of statistical tests related to Figure 2. Numerical data are provided in the Source Data File.

| # | Related to | Age | Descriptive statistics | N | Omnibus test | Age groups | Post-hoc tests | | |
|---|---|---|---|---|---|---|---|---|---|
| 1 | Figure 2C | | | | One-way ANOVA | | Tukey-Kramer test | | |
| | CaT frequency | P4 | 1.51 ± 0.16 min$^{-1}$ | 19 | F = 46 | P4 – P11 | q = 9.2 | P = 3.1 × $10^{-7}$ | df = 39 |
| | | P11 | 3.92 ± 0.40 min$^{-1}$ | 11 | P = 5.1 × $10^{-11}$ | P4 – P18 | q = 13 | P = 1.3 × $10^{-10}$ | df = 39 |
| | | P18 | 4.76 ± 0.29 min$^{-1}$ | 12 | | P11 – P18 | q = 2.9 | P = 0.11 | df = 39 |
| 2 | Figure 2E | | | | One-way ANOVA | | Tukey-Kramer test | | |
| | Gini coefficient of CaT frequencies | P4 | 0.35 ± 0.02 | 19 | F = 19 | P4 – P11 | q = 6.5 | P = 1.3 × $10^{-4}$ | df = 39 |
| | | P11 | 0.23 ± 0.02 | 11 | P = 1.8 × $10^{-6}$ | P4 – P18 | q = 2.9 | P = 0.11 | df = 39 |
| | | P18 | 0.41 ± 0.02 | 12 | | P11 – P18 | q = 8.5 | P = 1.5 × $10^{-6}$ | df = 39 |
| 3 | Figure 2G | | | | Kruskal-Wallis test | | Mann-Whitney test (Holm's method) | | |
| | CV2 of CaTs | P4 | 1.07 ± 0.01 | 19 | H = 27 | P4 – P11 | α = 0.025 | P = 5.6 × $10^{-5}$ | |
| | | P11 | 0.97 ± 0.01 | 11 | P = 1.8 × $10^{-6}$ | P4 – P18 | α = 0.017 | P = 2.0 × $10^{-6}$ | |
| | | P18 | 0.93 ± 0.01 | 12 | | P11 – P18 | α = 0.05 | P = 1.7 × $10^{-3}$ | |
| 4 | Results section | | | | Kruskal-Wallis test | | Mann-Whitney test (Holm's method) | | |
| | Gini coefficient of CV2 | P4 | 0.079 ± 0.005 | 19 | H = 15 | P4 – P11 | α = 0.017 | P = 1.5 × $10^{-4}$ | |
| | | P11 | 0.057 ± 0.004 | 11 | P = 6.2 × $10^{-4}$ | P4 – P18 | α = 0.05 | P = 0.59 | |
| | | P18 | 0.077 ± 0.003 | 12 | | P11 – P18 | α = 0.025 | P = 9.8 × $10^{-4}$ | |
| 5 | Results section | | | | One-way ANOVA | | | | |
| | $\Delta F/F_0$ noise | P4 | 0.101 ± 0.006 | 19 | F = 3 | P4 – P11 | | | |
| | | P11 | 0.126 ± 0.018 | 11 | P = 0.059 | P4 – P18 | | | |
| | | P18 | 0.122 ± 0.007 | 12 | | P11 – P18 | | | |

**Table S3.** Synopsis of statistical tests related to Figure 3. Numerical data are provided in the Source Data File.

| # | Related to | Age | Descriptive statistics | N | Omnibus test | Age groups | Post-hoc tests | | |
|---|---|---|---|---|---|---|---|---|---|
| 1 | Figure 3C | | | | Kruskal-Wallis test | | Mann-Whitney test (Holm's method) | | |
| | Time in continuous activity | P4 | 2.3 ± 0.7% | 19 | H = 30 | P4 – P11 | α = 0.025 | P = 9.9 × $10^{-6}$ | |
| | | P11 | 18.1 ± 4.8% | 11 | P = 3.6 × $10^{-7}$ | P4 – P18 | α = 0.017 | P = 5.7 × $10^{-8}$ | |
| | | P18 | 57 ± 9.4% | 12 | | P11 – P18 | α = 0.05 | P = 0.004 | |
| 2 | Figure 3E | | | | Kruskal-Wallis test | | Mann-Whitney test (Holm's method) | | |
| | Lomb-Scargle Bandpower (0.1–0.5 Hz) of Φ | P4 | (2.9 ± 0.4) × $10^{-4}$ | 19 | H = 20 | P4 – P11 | α = 0.017 | P = 9.9 × $10^{-6}$ | |
| | | P11 | (8.9 ± 1.8) × $10^{-4}$ | 11 | P = 4.5 × $10^{-5}$ | P4 – P18 | α = 0.05 | P = 0.33 | |
| | | P18 | (3.4 ± 0.3) × $10^{-4}$ | 12 | | P11 – P18 | α = 0.025 | P = 1.8 × $10^{-5}$ | |
| 3 | Figure 3F | | | | One-way ANOVA | | Tukey-Kramer test | | |
| | Network burst time fraction | P4 | 11.6 ± 0.8% | 19 | F = 34 | P4 – P11 | q = 11 | P = 3.2 × $10^{-9}$ | df = 39 |
| | | P11 | 22.6 ± 1.1% | 11 | P = 3.0 × $10^{-9}$ | P4 – P18 | q = 7.0 | P = 4.3 × $10^{-5}$ | df = 39 |
| | | P18 | 18.3 ± 1.2% | 12 | | P11 – P18 | q = 4.0 | P = 0.019 | df = 39 |
| 4 | Figure 3G | | | | Welch's ANOVA | | Games-Howell test | | |
| | Network burst duration | P4 | 0.66 ± 0.03 s | 19 | F = 39 | P4 – P11 | q = 0.79 | P = 0.84 | df = 20 |
| | | P11 | 0.63 ± 0.04 s | 11 | P = 8.4 × $10^{-8}$ | P4 – P18 | q = 11 | P = 8.4 × $10^{-8}$ | df = 25 |
| | | P18 | 0.38 ± 0.02 s | 12 | | P11 – P18 | q = 7.6 | P = 3.7 × $10^{-4}$ | df = 13 |
| 5 | Figure 3H | | | | Kruskal-Wallis test | | Mann-Whitney test (Holm's method) | | |
| | Network burst size | P4 | 7.3 ± 0.7% | 19 | H = 16 | P4 – P11 | α = 0.05 | P = 0.17 | |
| | | P11 | 8.9 ± 1.0% | 11 | P = 3.8 × $10^{-4}$ | P4 – P18 | α = 0.025 | P = 7.4 × $10^{-3}$ | |
| | | P18 | 4.8 ± 0.2% | 12 | | P11 – P18 | α = 0.017 | P = 1.5 × $10^{-6}$ | |
| 6 | Figure 3I | | | | Kruskal-Wallis test | | Mann-Whitney test (Holm's method) | | |
| | Mean participation rate of cells to NBs | P4 | 10.3 ± 0.8% | 19 | H = 9.4 | P4 – P11 | α = 0.025 | P = 0.008 | |
| | | P11 | 14.4 ± 1.2% | 11 | P = 0.009 | P4 – P18 | α = 0.05 | P = 0.39 | |
| | | P18 | 11.2 ± 0.4% | 12 | | P11 – P18 | α = 0.017 | P = 0.004 | |
| 7 | Results section | | | | Welch's ANOVA | | Games-Howell test | | |
| | Gini coefficient of participation rate of cells to NBs | P4 | 0.33 ± 0.02 | 19 | F = 25 | P4 – P11 | q = 7.9 | P = 1.7 × $10^{-5}$ | df = 28 |
| | | P11 | 0.21 ± 0.01 | 11 | P = 1.4 × $10^{-6}$ | P4 – P18 | q = 1.1 | P = 0.70 | df = 25 |
| | | P18 | 0.35 ± 0.02 | 12 | | P11 – P18 | q = 8.3 | P = 4.7 × $10^{-5}$ | df = 17 |

**Table S4.** Synopsis of statistical tests related to Figure 4. Numerical data are provided in the Source Data File.

| # | Related to | Age | Descriptive statistics | N | Omnibus test | Age groups | Post-hoc tests | | |
|---|---|---|---|---|---|---|---|---|---|
| 1 | Figure 4A | | | | One-way ANOVA | | Tukey-Kramer test | | |
| | Population coupling of all cells (σ = 3) | P4 | $(2.7 \pm 2.5) \times 10^{-4}$ | 19 | F = 16 | P4 – P11 | q = 7.7 | $P = 8.5 \times 10^{-6}$ | df = 39 |
| | | P11 | $(30.0 \pm 3.9) \times 10^{-4}$ | 11 | $P = 9.0 \times 10^{-6}$ | P4 – P18 | q = 1.1 | P = 0.72 | df = 39 |
| | | P18 | $(6.6 \pm 4.7) \times 10^{-4}$ | 12 | | P11 – P18 | q = 6.0 | $P = 3.4 \times 10^{-4}$ | df = 39 |
| 2 | Figure 4B | | | | Welch's ANOVA | | Games-Howell test | | |
| | Fraction of cells with significant population coupling (σ = 3) | P4 | 8.5 ± 0.7% | 19 | F = 8.7 | P4 – P11 | q = 5.8 | $P = 3.8 \times 10^{-3}$ | df = 12 |
| | | P11 | 17.6 ± 2.1% | 11 | $P = 2.4 \times 10^{-3}$ | P4 – P18 | q = 2.4 | P = 0.23 | df = 15 |
| | | P18 | 11.7 ± 1.7% | 12 | | P11 – P18 | q = 3.1 | P = 0.10 | df = 20 |
| 3 | Figure 4C | | | | Welch's ANOVA | | | | |
| | Population coupling of significant cells (dt = 3) | P4 | 0.021 ± 0.002 | 19 | F = 1.2 | | | | |
| | | P11 | 0.018 ± 0.001 | 11 | P = 0.32 | | | | |
| | | P18 | 0.019 ± 0.001 | 12 | | | | | |
| 4 | Figure 4E | | | | Kruskal-Wallis test | | Mann-Whitney test (Holm's method) | | |
| | Fraction of significant pairs for STTC (dt = 3) | P4 | 13.3 ± 1.8% | 19 | H = 18 | P4 – P11 | α = 0.05 | P = 0.07 | |
| | | P11 | 23.5 ± 4.8% | 11 | $P = 1.2 \times 10^{-4}$ | P4 – P18 | α = 0.025 | $P = 1.4 \times 10^{-3}$ | |
| | | P18 | 5.3 ± 0.7% | 12 | | P11 – P18 | α = 0.017 | $P = 5.9 \times 10^{-6}$ | |
| 5 | Figure 4G | | | | Welch's ANOVA | | Games-Howell test | | |
| | STTC of significant pairs (dt = 3) | P4 | 0.183 ± 0.014 | 19 | F = 20 | P4 – P11 | q = 8.9 | $P = 5.5 \times 10^{-6}$ | df = 23 |
| | | P11 | 0.087 ± 0.006 | 11 | $P = 9.4 \times 10^{-6}$ | P4 – P18 | q = 8.5 | $P = 1.8 \times 10^{-5}$ | df = 20 |
| | | P18 | 0.095 ± 0.004 | 12 | | P11 – P18 | q = 1.6 | P = 0.50 | df = 17 |
| 6 | Figure 4H | | | | | | | | |
| | Spearman rho of STTC (dt = 3) to neuron-neuron distance | P4 | -0.046 | 12394 | $P = 3.2 \times 10^{-7}$ | | | | |
| | | P11 | -0.032 | 17296 | $P = 2.1 \times 10^{-5}$ | | | | |
| | | P18 | 0.016 | 4400 | P = 0.27 | | | | |

**Table S5.** Synopsis of statistical tests related to Figure 5. Numerical data are provided in the Source Data File.

| # | Related to | Age | Descriptive statistics | N | Omnibus test | Age groups | Post-hoc tests | |
|---|---|---|---|---|---|---|---|---|
| 1 | Figure 5B | | | | Kruskal-Wallis test | | Mann-Whitney test (Holm's method) | |
| | Gobal similarity of activity patterns (Win = 10) | P4 | 0.055 ± 0.010 | 19 | H = 17 | P4 – P11 | α = 0.025 | P = 2.4 × $10^{-4}$ |
| | | P11 | 0.017 ± 0.002 | 11 | P = 1.7 × $10^{-4}$ | P4 – P18 | α = 0.05 | P = 0.39 |
| | | P18 | 0.055 ± 0.007 | 12 | | P11 – P18 | α = 0.017 | P = 1.8 × $10^{-5}$ |
| 2 | Figure 5C | | | | Kruskal-Wallis test | | Mann-Whitney test (Holm's method) | |
| | Number of motifs (Win = 10) | P4 | 5.9 ± 0.4 | 19 | H = 12 | P4 – P11 | α = 0.017 | P = 9.8 × $10^{-4}$ |
| | | P11 | 2.5 ± 0.9 | 11 | P = 3.0 × $10^{-3}$ | P4 – P18 | α = 0.05 | P = 0.41 |
| | | P18 | 7.0 ± 1.1 | 12 | | P11 – P18 | α = 0.025 | P = 5.6 × $10^{-3}$ |
| 3 | Figure 5D | | | | Kruskal-Wallis test | | Mann-Whitney test (Holm's method) | |
| | Fraction of patterns in motifs (Win = 10) | P4 | 17.8 ± 1.2% | 19 | H = 15 | P4 – P11 | α = 0.05 | P = 0.09 |
| | | P11 | 16.6 ± 5.9% | 11 | P = 4.8 × $10^{-4}$ | P4 – P18 | α = 0.017 | P = 9.5 × $10^{-5}$ |
| | | P18 | 32.4 ± 4.7% | 12 | | P11 – P18 | α = 0.025 | P = 4.5 × $10^{-3}$ |

**Table S6.** Synopsis of statistical tests related to Figure 6. Numerical data are provided in the Source Data File.

| # | Related to | Group | Descriptive statistics | N | Statistics | | |
|---|---|---|---|---|---|---|---|
| 1 | Figure 6B | | | | T-Test for paired samples | | |
| | Respiration rate | $N_2O$ | 184 ± 7 $min^{-1}$ | 12 | t = 0.50 | df = 11 | P = 0.62 |
| | | Unanesth. | 188 ± 6 $min^{-1}$ | | | | |
| 2 | Figure 6C | | | | T-Test for paired samples | | |
| | Heartbeat | $N_2O$ | 353 ± 14 $min^{-1}$ | 12 | t = 1.12 | df = 11 | P = 0.29 |
| | | Unanesth. | 338 ± 9 $min^{-1}$ | | | | |
| 3 | Figure 6D | | | | T-Test for paired samples | | |
| | Movement periods Total time | $N_2O$ | 3.8 ± 0.6% | 12 | t = 5.7 | df = 11 | P = $1.4 \times 10^{-4}$ |
| | | Unanesth. | 17.1 ± 2.3% | | | | |
| 4 | Results section | | | | T-Test for paired samples | | |
| | Movement periods Occurence frequency | $N_2O$ | 1.9 ± 0.2 $min^{-1}$ | 12 | t = 4.7 | df = 11 | P = $6.1 \times 10^{-4}$ |
| | | Unanesth. | 5.1 ± 0.7 $min^{-1}$ | | | | |
| 5 | Results section | | | | T-Test for paired samples | | |
| | Movement periods Duration | $N_2O$ | 1.2 ± 0.1 s | 12 | t = 4.4 | df = 11 | P = 0.001 |
| | | Unanesth. | 2.1 ± 0.2 s | | | | |
| 6 | Figure 6G | | | | T-Test for paired samples | | |
| | Network burst time fraction | $N_2O$ | 20.4 ± 1.5% | 12 | t = 0.84 | df = 11 | P = 0.42 |
| | | Unanesth. | 21.9 ± 1.3% | | | | |
| 7 | Figure 6H | | | | T-Test for paired samples | | |
| | Fraction of significant pairs for STTC (dt = 3) | $N_2O$ | 8.7 ± 1.1% | 12 | t = 3.1 | df = 11 | P = 0.01 |
| | | Unanesth. | 6.0 ± 0.8% | | | | |
| 8 | Figure S3C | | | | T-Test for paired samples | | |
| | CaT frequency | $N_2O$ | 2.52 ± 0.25 $min^{-1}$ | 12 | t = 2.4 | df = 11 | P = 0.035 |
| | | Unanesth. | 3.13 ± 0.21 $min^{-1}$ | | | | |
| 9 | Figure S3D | | | | T-Test for paired samples | | |
| | Lomb-Scargle Band-power (0.1–0.5 Hz) of Φ | $N_2O$ | $(3.7 \pm 0.4) \times 10^{-4}$ | 12 | t = 0.57 | df = 11 | P = 0.58 |
| | | Unanesth. | $(3.4 \pm 0.5) \times 10^{-4}$ | | | | |
| 10 | Figure S3E | | | | T-Test for paired samples | | |
| | Network burst size | $N_2O$ | 5.7 ± 0.4% | 12 | t = 1.3 | df = 11 | P = 0.22 |
| | | Unanesth. | 5.1 ± 0.5% | | | | |
| 11 | Figure S3F | | | | T-Test for paired samples | | |
| | STTC of significant pairs (dt = 3) | $N_2O$ | 0.104 ± 0.003 | 12 | t = 0.45 | df = 11 | P = 0.66 |
| | | Unanesth. | 0.103 ± 0.004 | | | | |

**Table S6** (continued)

| | | | | | | | |
|---|---|---|---|---|---|---|---|
| 12 | Results section | | | | Wilcoxon signed-rank test (exact) | | |
| | Population coupling of all cells (σ = 3) | $N_2O$ | (16.2 ± 2.8) × $10^{-4}$ | 12 | W = 76 | z = 2.9 | P = 0.001 |
| | | Unanesth. | (-2.5 ± 4.5) × $10^{-4}$ | | | | |
| 13 | Figure S3G | | | | T-Test for paired samples | | |
| | Fraction of cells with sign. PC (σ = 3) | $N_2O$ | 9.8 ± 0.8% | 12 | t = 2.9 | df = 11 | P = 0.01 |
| | | Unanesth. | 7.3 ± 0.7% | | | | |
| 14 | Results section | | | | T-Test for paired samples | | |
| | PC of significant cells (dt = 3) | $N_2O$ | 0.019 ± 0.001 | 12 | t = 0.33 | df = 11 | P = 0.74 |
| | | Unanesth. | 0.019 ± 0.001 | | | | |
